# *Clostridioides difficile* spore-entry into intestinal epithelial cells contributes to recurrence of the disease

**DOI:** 10.1101/2020.09.11.291104

**Authors:** Pablo Castro-Córdova, Paola Mora-Uribe, Rodrigo Reyes-Ramírez, Glenda Cofré-Araneda, Josué Orozco-Aguilar, Christian Brito-Silva, María José Mendoza-León, Sarah A. Kuehne, Nigel P. Minton, Marjorie Pizarro-Guajardo, Daniel Paredes-Sabja

## Abstract

*Clostridioides difficile* spores produced during infection are essential for the recurrence of the disease. However, how *C. difficile* spores persist in the intestinal mucosa to cause recurrent infection remains unknown. Here, we show that *C. difficile* spores gain entry into the intestinal mucosa via fibronectin-α_5_β_1_ and vitronectin-α_v_β_1_ specific-pathways. The spore-surface exosporium BclA3 protein is essential for both spore-entry pathways into intestinal epithelial cells. Furthermore, *C. difficile* spores of a *bclA3* isogenic mutant exhibited reduced entry into the intestinal mucosa and reduced recurrence of the disease in a mouse model of the disease. Inhibition of *C. difficile* spore-entry led to reduced spore-entry into the intestinal epithelial barrier and recurrence of *C. difficile* infection *in vivo*. These findings suggest that *C. difficile* spore-entry into the intestinal barrier is a novel mechanism of spore-persistence that can contribute to infection recurrence and have implications for the rational design of therapies.

## Introduction

*Clostridioides difficile* is a strict anaerobic Gram-positive pathogenic bacterium that forms highly resistant spores that easily persist in the environment and contribute to transmission of *C. difficile* infections (CDI) through fecal-oral route^1^. Disruption of the gut microbiota by the use of broad-spectrum antibiotics leads to an optimal environment for *C. difficile* colonization and proliferation in the colon and disease manifestation. CDI currently leads hospital-acquired diarrhea associated to antibiotics in United States and world-wide^2^. In the US alone, ∼500,000 patients per year become infected with CDI, and mortality rates reach ∼8% of total patients^2^. The annual cost of CDI to the health care system is estimated in ∼US 4.8 billion^2^. Treatment of CDI usually involves antibiotic therapy, typically vancomycin or metronidazole and, most recently, fidaxomicin^2^, which, although resolves the infection in ∼95% of the cases, leads to recurrence of CDI (R-CDI) in 15-30% of the individuals^3-5^.

During infection, *C. difficile* produces two major virulence factors, toxins TcdA and TcdB, which are responsible for the clinical manifestation of the disease, induce pro-inflammatory cytokines, disruption of tight junctions, detachment of intestinal epithelial cells (IEC) and loss of trans-epithelial barrier^6^. *C. difficile* also initiates a sporulation pathway that leads to the production of new metabolically dormant spores in the host’s intestine^1^. *In vivo*, spore-formation is essential for the recurrence of the disease^7^. Moreover, spore-based therapies that remove *C. difficile* spores from the intestinal mucosa contribute to reduce recurrence of the disease in animal models^8^.

Recent *in vivo* studies in the laboratory strain 630 suggest that the spore surface mucus-binding protein, peroxiredoxin-chitinase CotE, and the exosporium collagen-like BclA1 proteins are required for the colonization and infectivity in a mouse model of CDI^9,10^. However, the surface layer of 630 spores does not resemble that of clinically relevant strains which exhibit hair-like projections in their spore surface; structures that are absent in strain 630^1,11,12^. Importantly, most clinically relevant sequenced *C. difficile* isolates, including isolates of the epidemically relevant 027 ribotype, have a truncated *bclA1* due to a premature stop codon in the N-terminal domain^13^, resulting in the translation of a small polypeptide, which localizes to the spore surface^10^; thus limiting the breadth and depth of these results.

*C. difficile* spores exhibit high levels of adherence to intestinal epithelial cells (IECs) *in vitro*^14,15^, and that the hair-like projections of *C. difficile* spores come in close proximity with the microvilli of differentiated Caco-2 cells; furthermore, *C. difficile* spores interact in a dose-dependent manner with fibronectin (Fn) and vitronectin (Vn)^15^, two extracellular matrix proteins used by several enteric pathogens to infect the host^16,17^. However, the mechanisms that underline how these interactions contribute to *C. difficile* spore-persistence *in vivo* and contribute to the recurrence of the disease remain unclear.

Herein we first demonstrate that *C. difficile* spores gain entry into the intestinal epithelial barrier of mice and that spore-entry into IECs requires serum-molecules, specifically Fn and Vn, that are luminary accessible in the colonic mucosa. We also demonstrate that the spore-entry pathway into IECs is Fn-α_5_β_1_ and Vn-α_v_β_1_. Next, we demonstrate that the spore surface collagen-like BclA3 protein is essential for spore-entry into IECs through these pathways *in vitro* and essential for spore adherence to the intestinal mucosa. Importantly, BclA3 contributes to the recurrence of the disease in mice. We also observed the therapeutic potential of blocking spore-entry into the intestinal epithelial barrier and how co-administration of nystatin with vancomycin reduces spore-persistence and R-CDI in mice. Together, our results reveal a novel mechanism employed by *C. difficile* spores that contributes to R-CDI, which involves gaining intracellular access into the intestinal barrier via BclA3-Fn-α_5_β_1_ and BclA3-Vn-α_v_β_1_ specific, and that blocking spore-entry contributes to reduced recurrence of the disease.

## Results

### *C. difficile* spores internalize into the intestinal barrier *in vivo*

To study the interaction of *C. difficile* spores and the host’s intestinal barrier, we used a colonic/ileal loop assay infected with *C. difficile* spores for 5 hours^18^, where *C. difficile* R20291 spores, were labeled with anti-spore antibodies^13,18^. We observed similar levels of adherence of *C. difficile* spores to the colonic and ileum mucosa (Fig. 1 a–c), with no preference for the site of spore-adherence in both colonic and ileum mucosa (Fig. 1d, Extended Data Fig. 1 and 2). Strikingly, we observed that *C. difficile* spores were able to cross the mucosal barrier in the colonic/ileal loop assay (Fig. 1a, b, e, f, Supplementary Video 1 and 2 and Extended Data Fig. 3, and 4). We observed that 4.6 and 3.7 spores per 10^5^ μm^−2^ were able to cross the mucosal barrier in colonic and ileal loops (Fig. 1e), corresponding to 0.92% ± 0.30% and 1.04%±0.48 of the total spores, respectively. In the colonic mucosa internalized *C. difficile* spores were found to homogeneously localize 10 to 30 μm from the colonic surface and 5 to 50 μm from the closest crypt membrane, while in the ileum mucosa spores were homogeneously found at 15 to 70 μm from the villus tip and 10 to 50 μm from the villus membrane in ileal loops, (Extended Data Fig. 5), indicating multiple sites of entry in colon and ileum.

**Fig. 1.**
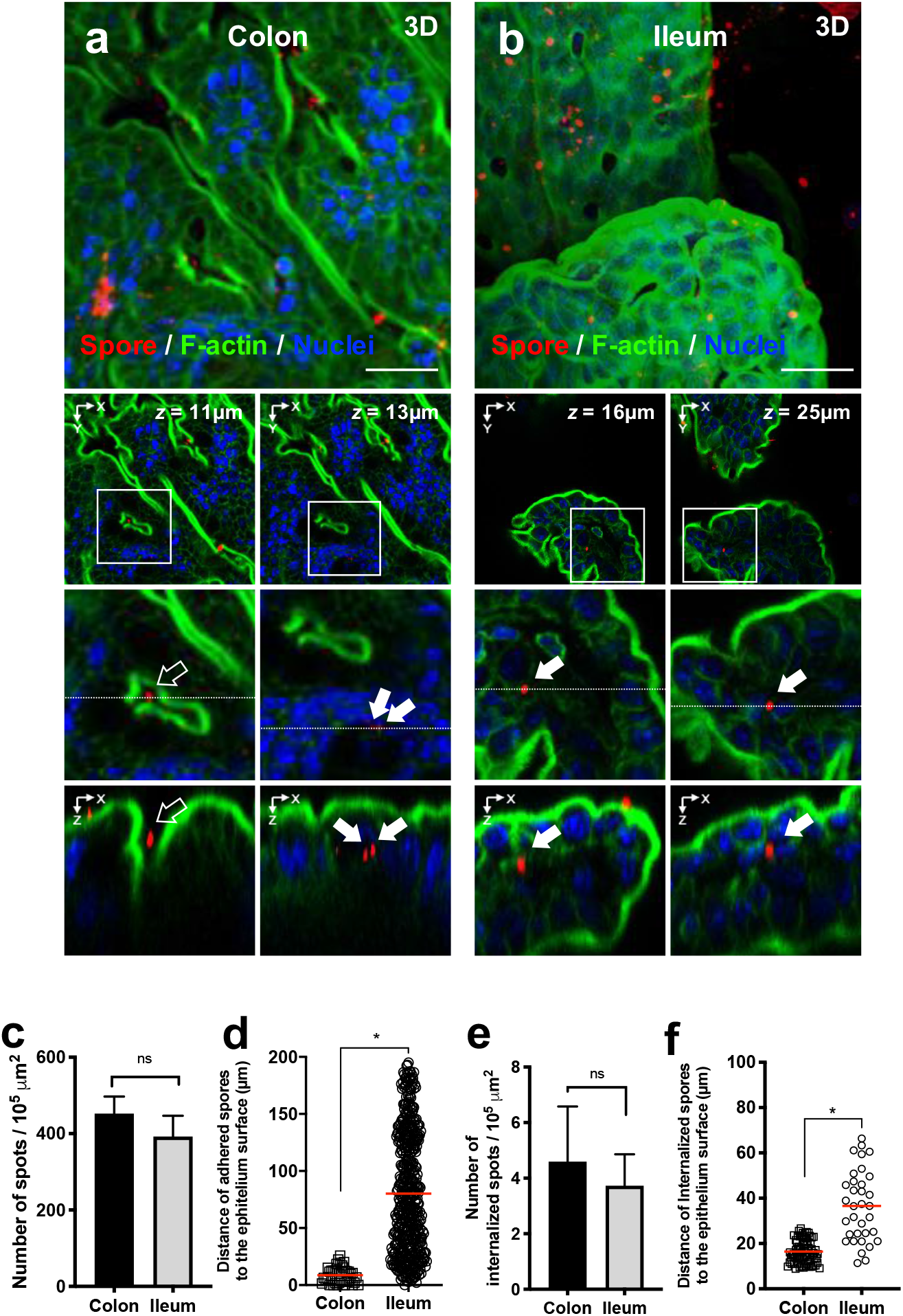
*C. difficile* adherence and internalization into intestinal barrier *in vivo*. **a, b** Confocal micrographs of fixed whole-mount of **a** colonic mucosa, and **b** ileum mucosa (SI) loop of C57BL/6 mice infected with 5 × 10^8^ *C. difficile* R20291 spores for 5 h. *C. difficile* spores are shown in red, F-actin is shown in green and nuclei in blue (fluorophores colors were digitally reassigned for a better representation). **c**, Adherence of *C. difficile* spores to SI and colonic mice tissue. **d**, Distance of adhered spores from the villus tip or from the colonic epithelial apical surface. **e**, Quantification of internalized *C. difficile* spores in the SI and colon. **f**, Distance of internalized spores from the villus tip for the ileum or from the epithelium surface for the colon. Micrographs are representative of mice (*n* = 3). White arrow indicates internalized *C. difficile* spores, empty arrow indicates adhered *C. difficile* spore. Scale bar, 20 µm. Error bars indicate the mean ± S.E.M. Statistical analysis was performed by Mann-Whitney test, ns indicates non-significant differences, * *P* < 0.0001.

**Fig. 2.**
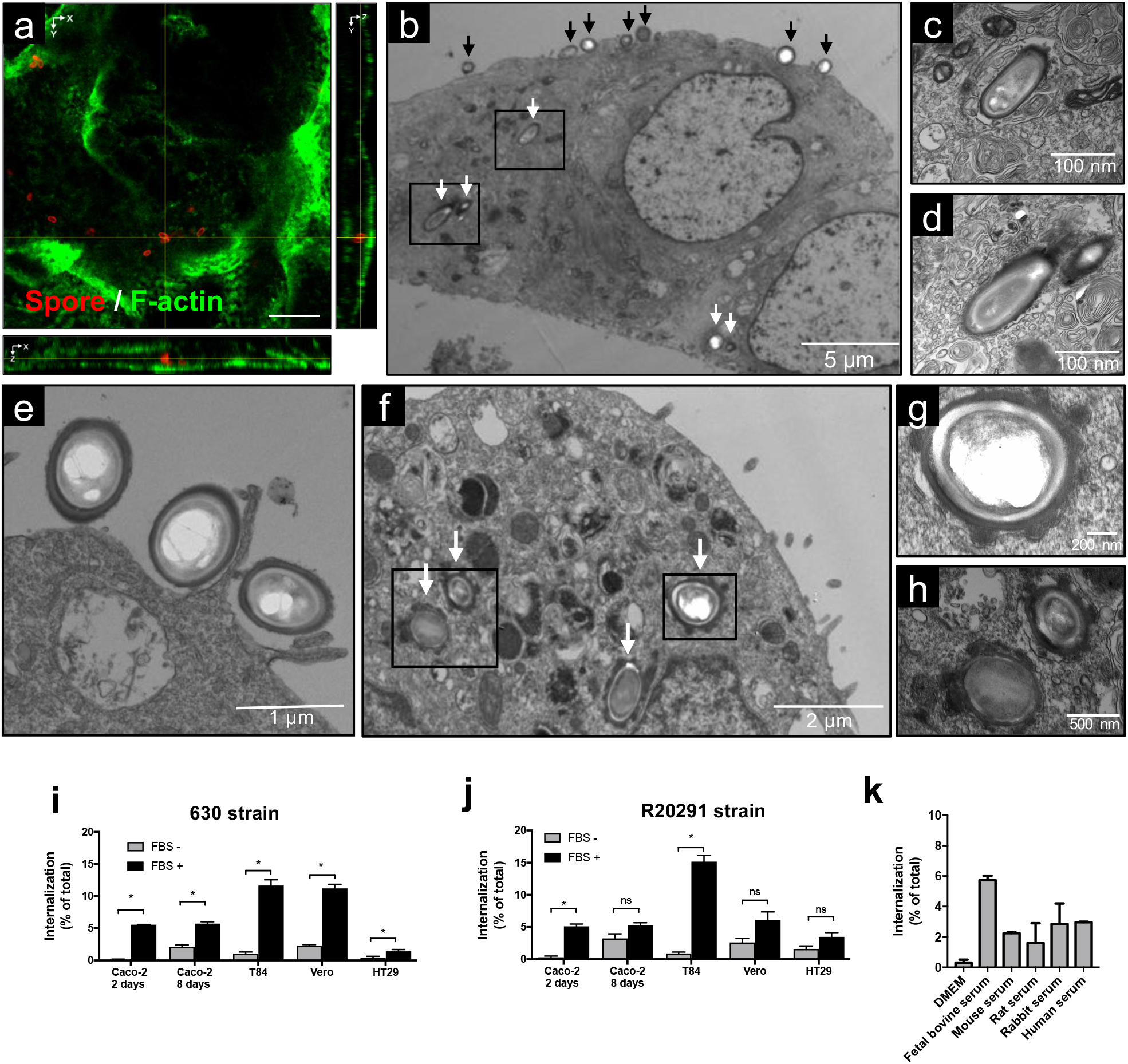
*C. difficile* spores are internalized by intestinal epithelial cells. **a** Confocal microscopy of an internalized *C. difficile* 630 spore in T84 cells. *C. difficile* spores are shown in red, F-actin is shown in green (fluorophores colors were digitally reassigned for a better representation). Yellow lines indicate an internalized spore. **b**−**e** TEM of differentiated T84 cell monolayers infected with *C. difficile* 630 spores. Black and white arrows denote extracellular and intracellular *C. difficile* spores, respectively. **c, d** are magnifications of black squares of **b. e** show an adhered *C. difficile* spore and an apical membrane extension of T84 cells surrounding *C. difficile* spores. **f**−**h** TEM of differentiated monolayers of Caco-2 cells infected with *C. difficile* R20291 spores. White arrows in panel **f** indicate internalized *C. difficile* spores. **g, h** are magnifications of black boxes in panel **f**. Internalization of *C. difficile* spores **i** strain 630 and, **j** R20291 pre-incubated with FBS or culture media in Caco-2 undifferentiated (2 days), differentiated (8 days), T84, Vero, and HT29. **k**, Internalization of *C. difficile* spores pre-incubated with serum of different mammalian species in Caco-2 cells. Error bars indicate the mean ± S.E.M. Scale bars **a** 5 μm; **c, d**, 100 nm; **e**, 1 μm; **f**, 2 μm; **g**, 200nm; **h**, 500nm.

**Fig. 3.**
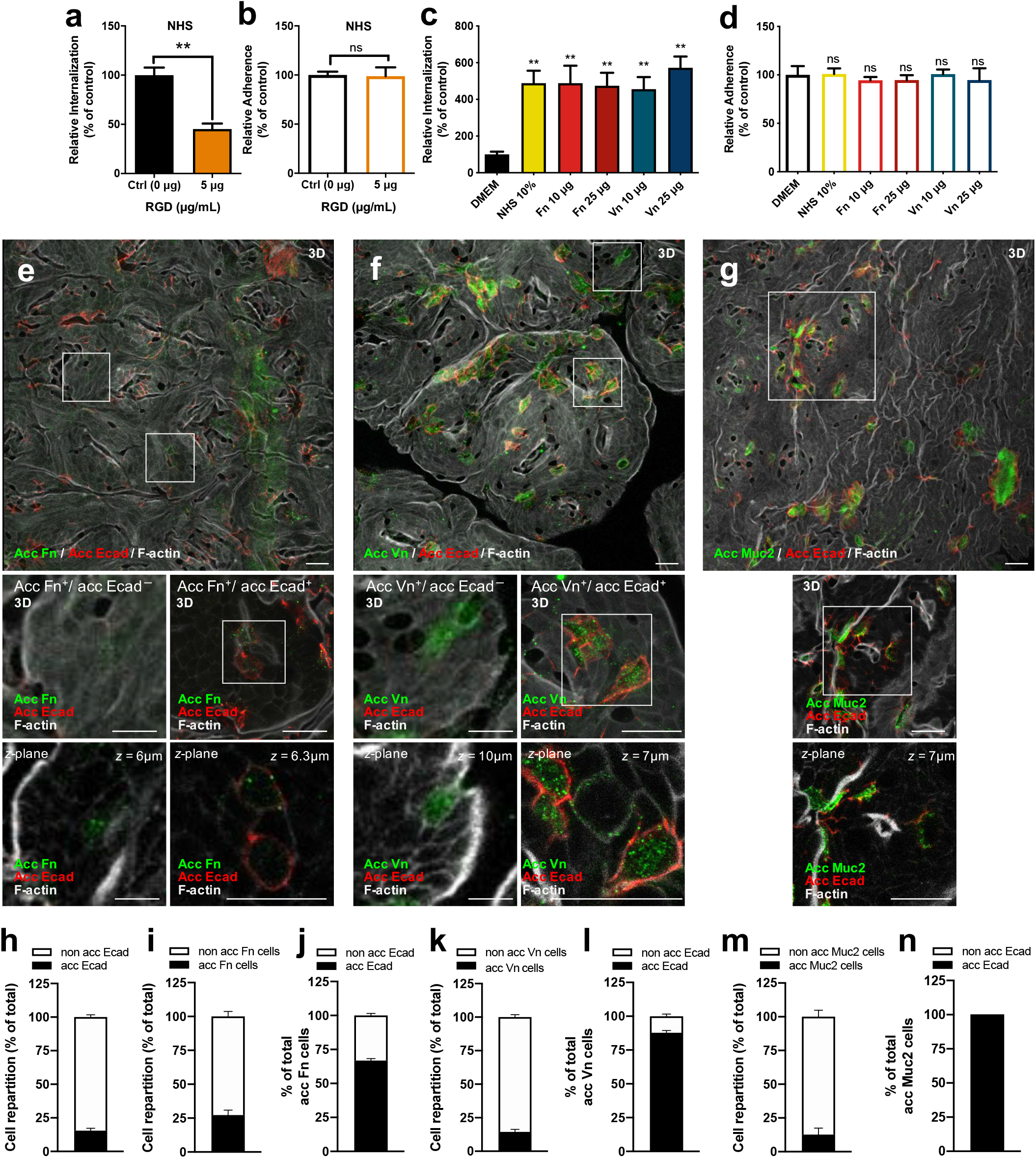
*C. difficile* spore internalization requires Fn and Vn which are luminally accessible in the intestinal barrier. **a** Internalization **b** and adherence of *C. difficile* spores pre-incubated with NHS in differentiated Caco-2 monolayers in the presence of 5 μg RGD peptide. **c, d** differentiated Caco-2 cells were infected with *C. difficile* spores pre-incubated with DMEM, NHS, 10, or 20 μg mL^−1^ Fn or Vn. Data shown in each panel are normalized to the control (0 µg mL^−1^ RGD or DMEM) and represent the mean of three independent experiments. **e, f, g** confocal micrographs of fixed whole-mount of the healthy colon of C57BL/6 mice for **e**, acc Fn; **f**, acc Vn; and **g**, acc Muc2 with acc Ecad. The main figure shown a 3D projection, below magnifications and a z-stack of representative cells with different immunostaining. **h** Shown the cell repartition of cell immunodetected for acc Ecad. **i** Shown the cell repartition of cells immunodetected for acc Fn. **j** Cell repartition of total acc Fn cells that were immunodetected for acc E-cad. **k** shown the cell repartition of cells immunodetected for acc Vn. **l** Cell repartition of total acc Vn cells that were immunodetected for acc E-cad. **m** shown the cell repartition of cells immunodetected for acc Muc2. **n** Cell repartition of total acc Muc2 cells that were immunodetected for acc E-cad. Acc-FN, acc-VN and Muc2 is shown in green, Acc-Ecad is shown in red and F-actin in grey (fluorophores colors were digitally reassigned for a better representation). Scale bar, 20µm. Micrographs are representative of mice (*n* = 2). 1,000 - 1,200 cells were counted for each mouse in an area 84,628µm^2^. Error bars indicate mean ± S.E.M. Statistical analysis was performed by Student’s *t*-test, ns indicates non-significant differences, * *P* < 0.05 and ** *P* < 0.001. Scale bar, 20 µm.

**Fig. 4.**
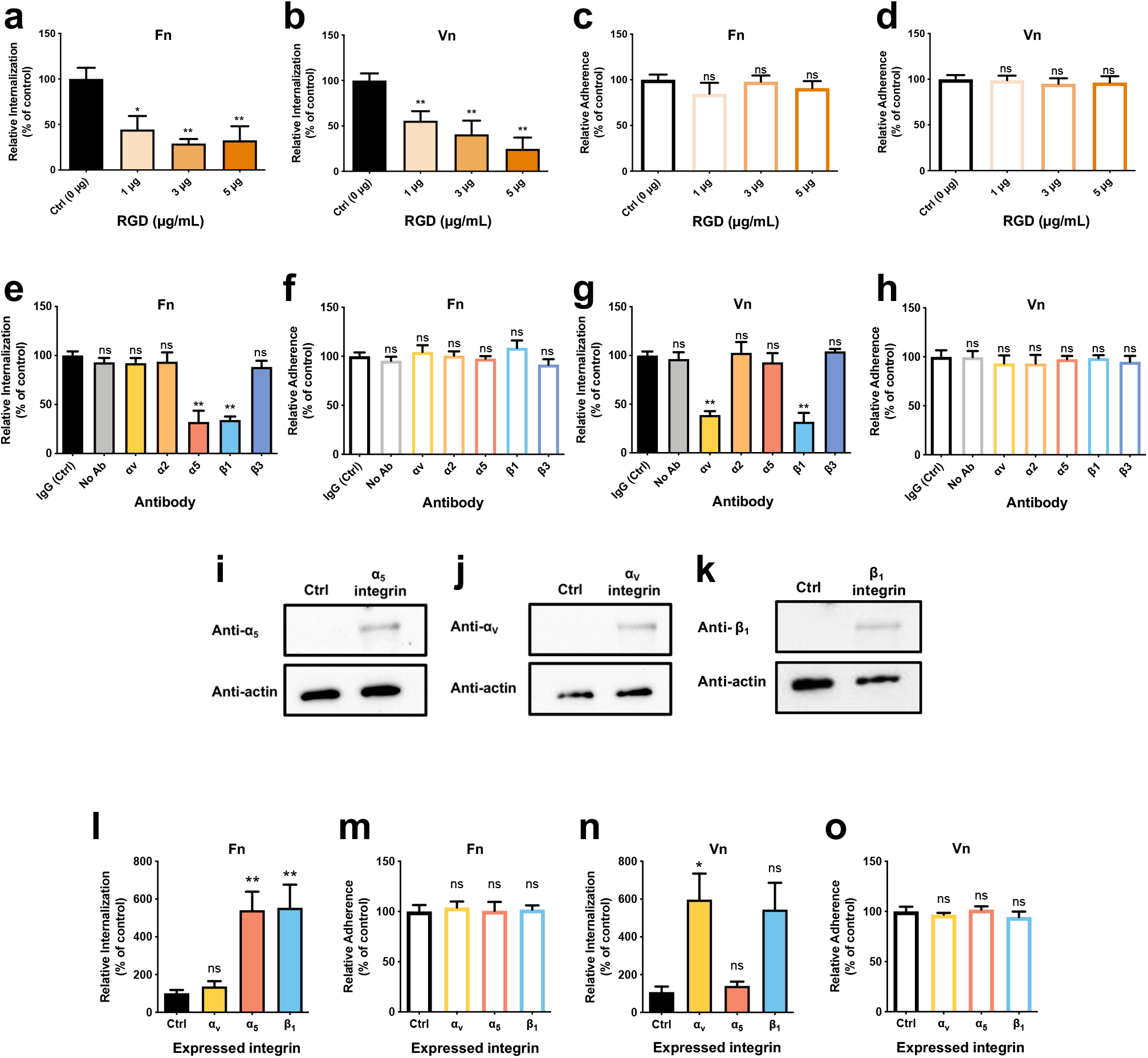
*C. difficile* spore internalization via fibronectin-α_5_β_1_ and vitronectin-α_v_β_1_ integrins intestinal epithelial cells. *C. difficile* spore internalization in **a**−**d** differentiated Caco-2 cells incubated for 1 h with 1, 3, and 5 µg/mL of RGD peptide and infected with *C. difficile* spores pre-incubated for 1 h with **a, c**, 10 μg mL^−1^ Fn and **b, d**, 10 μg mL^−1^ Vn. **e**−**h** Differentiated Caco-2 monolayers were incubated for 1 h with 10 µg mL^−1^ of antibody against α_v_, α_2,_ α_5_, β_1_, β_3_, non-immune IgG antibody or without antibody. Then were infected with *C. difficile* spores R20291 pre-incubated with 10 μg mL^−1^ of **e, f**, Fn or **g, h**, Vn. **i**−**k**, show immunoblotting of cell lysates of CHO cells transfected with ectopic expression of **i** α_5_ (∼120 kDa); **j** α_v_ (∼120 kDa); and **k** β_1_ (∼120 kDa) and alpha-tubulin as a loading control (50 kDa). *C. difficile* spore **l, n** internalization or **m, o** adherence in CHO cells ectopically expressing α_v_, α_5_, β_1_ integrins, of spores pre-treated 1 h with **l, o** 10 μg mL^−1^ of Fn and **n, o**, 10 μg mL^−1^ of Vn. Data shown in each panel are normalized to the control (0 µg mL^−1^ RGD, DMEM, or IgG) and represent the mean of three independent experiments. Error bars indicate mean ± S.E.M. Statistical analysis was performed by Student’s *t*-test, ns indicates non-significant differences, * *P* < 0.05 and ** *P* < 0.001.

**Fig. 5.**
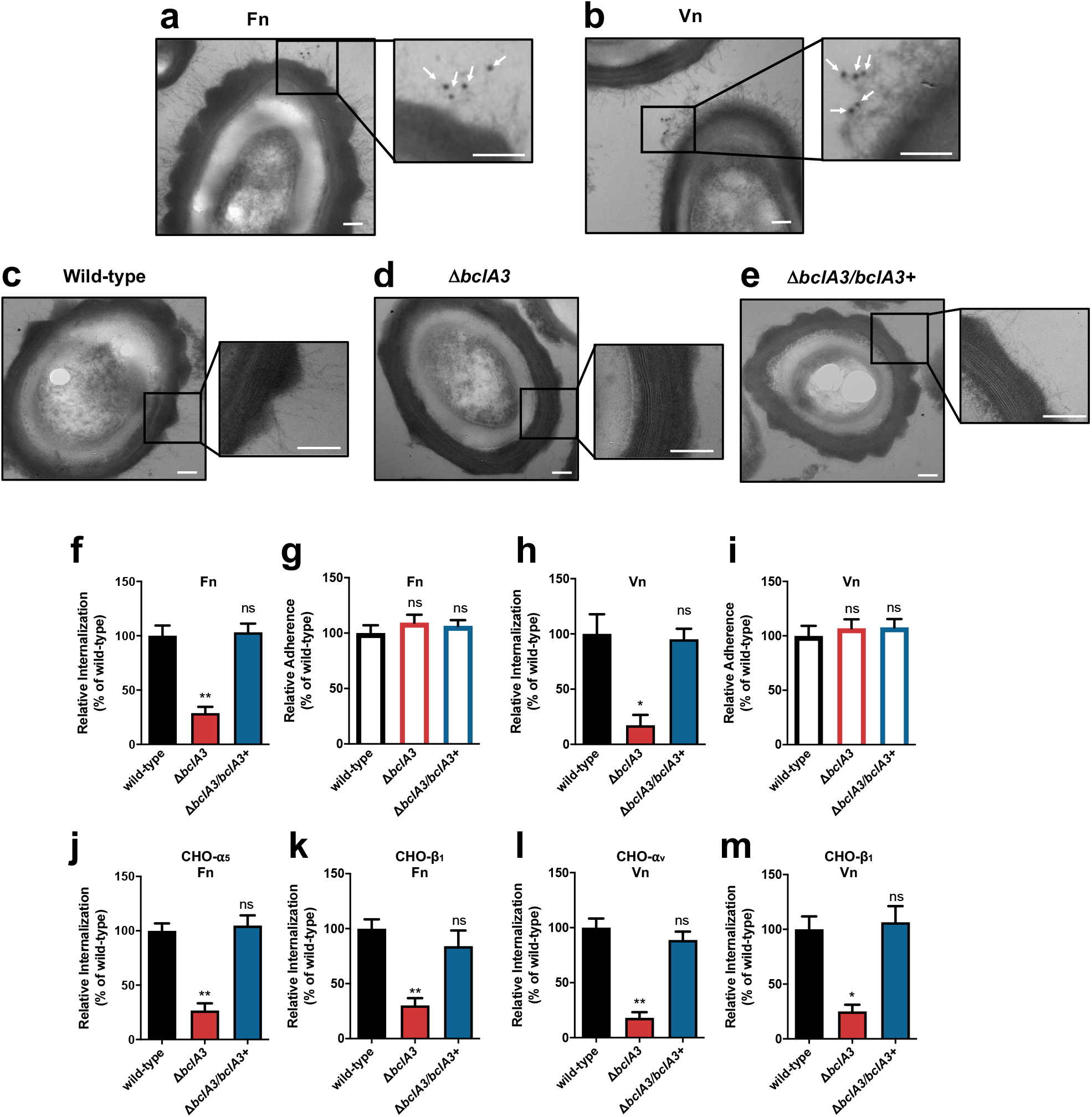
The collagen-like exosporium protein BclA3 is required for spore-entry into intestinal epithelial cells via Fn-α_5_β_1_ and Vn-α_v_β_1_. **a, b** Immunogold of Fn and Vn binding to the hair-like extensions of *C. difficile* spores. *C. difficile* spores R20291 were incubated with 10 µg mL^−1^ Fn or Vn for 1 h. Samples were processed and visualized for TEM. White arrows indicate anti-Fn or -Vn rabbit antibody and anti–rabbit–gold 12 nm antibody complex. **ce** TEM of wild-type (Δ*pyrE*/*pyrE*^+^), Δ*bclA3* and Δ*bclA3/bclA3+ C. difficile* R20291 spores. **f, h** Internalization and **g, i** adherence in differentiated Caco-2 cells of wild-type (Δ*pyrE*/*pyrE*^+^), Δ*bclA3* and Δ*bclA3/bclA3+ C. difficile* R20291 spores pre-incubated with **f, g** 10 μg mL^−1^ Fn and **h, i** 10 μg mL^−1^ Vn. **j, l** internalization and **k, m** adherence in CHO cells ectopically expressing: **j** α_5_; **k, m**; β_1_ or **l** α_v_ infected with wild-type (Δ*pyrE*/*pyrE*^+^), Δ*bclA3* and Δ*bclA3/bclA3+ C. difficile* R20291 spores pre-incubated with **j, k** 10 μg mL^−1^ Fn and **l, m** 10 μg mL^−1^ Vn. Data shows internalization and adherence normalized to wild-type spores and represented the mean of three independent experiments. Error bars indicate the mean ± S.E.M. Statistical analysis was performed by Student’s *t*-test, ns indicates non-significant differences, * *P* < 0.005, \*\**P* < 0.0001. **a**−**e**, Scale bar, 100 nm.

### *C. difficile* spore-entry into intestinal epithelial cells requires serum components *in vitro*

Our previous *in vitro* studies in IECs were conducted in the absence of fetal bovine serum (FBS) and did not evidence internalized spores^14,15,18,19^. Therefore, we assess if FBS contributed to spore-entry by confocal fluorescence microscopy by analyzing monolayers of polarized T84 IECs (Fig. 2a and Extended Data Fig. 6a, b) and differentiated Caco-2 cells (Extended Data Fig. 6c, d) which were infected with *C. difficile* spores of the epidemically relevant R20291 and the commonly used strain 630 in the presence of FBS. In both cell lines, several intracellular spores of strain *C. difficile* 630 were found to be located between the apical and basal actin cytoskeleton (Fig. 2a and Extended Data Fig 6). To obtain convincing evidence of entry of *C. difficile* spores into IECs, we analyzed polarized monolayers of T84 and Caco-2 cell lines infected with *C. difficile* 630 or R20291 spores using transmission electron microscopy (TEM). Electron micrographs evidence that some *C. difficile* spores were found extracellularly in the apical membrane, while others were found intracellularly (Fig. 2b–d). Intracellular *C. difficile* spores were surrounded by an endosomal-like membrane (Fig. 2c, d). Notably, the formation of membrane lamellipodia-like protrusions and circular ruffle surrounding *C. difficile* 630 spores were evidenced at the site of attachment of *C. difficile* spores to the apical membrane (Fig. 2e), suggesting macropinocytosis-like endocytosis of *C. difficile* spores. Intracellular spores of the strain, R20291, were also evidenced in differentiated Caco-2 cells (Fig. 2f–h).

**Fig. 6.**
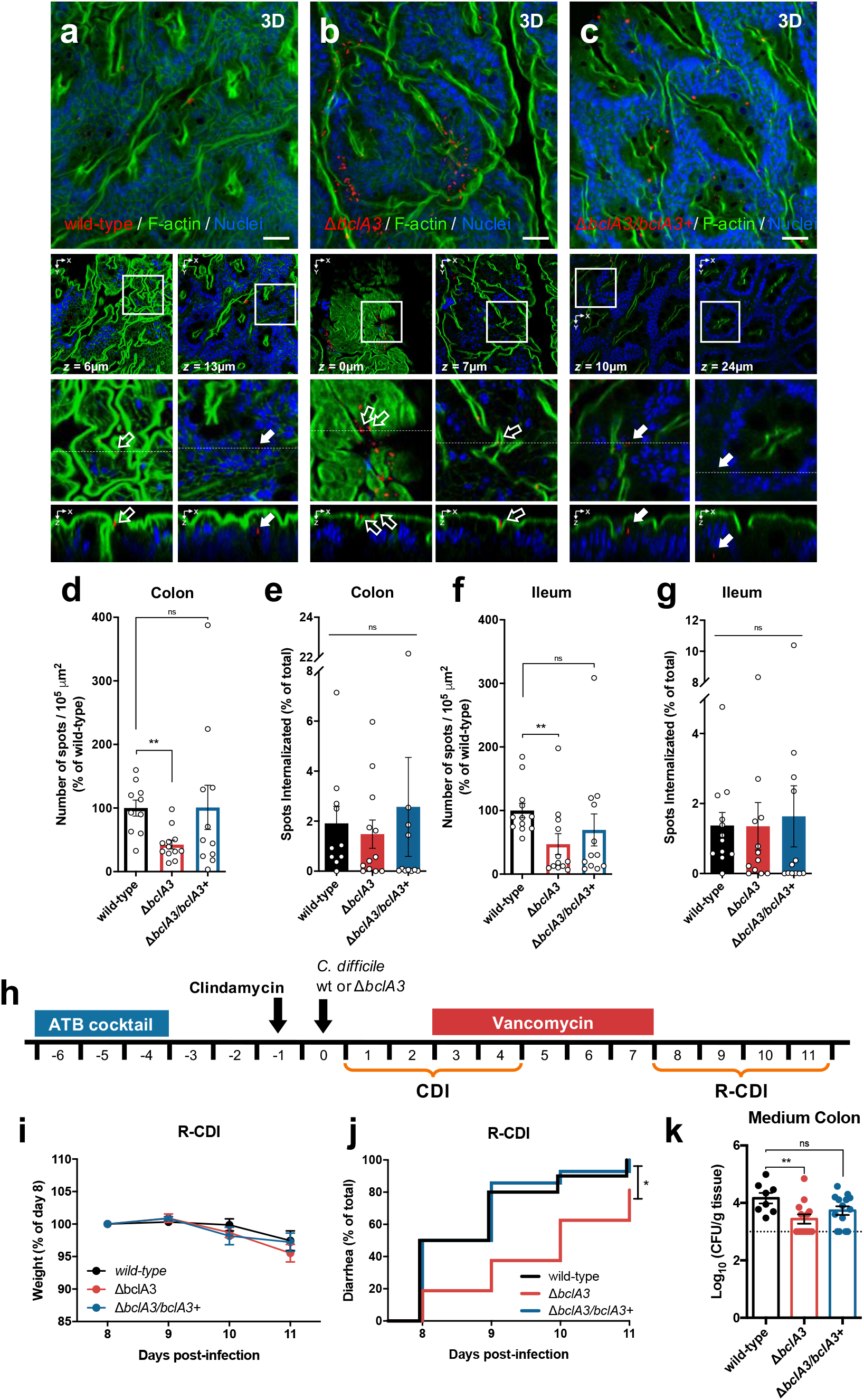
BclA3 is involved in *C. difficile* spore adherence to the intestinal mucosa and delays the onset of diarrhea during R-CDI. Intestinal loops of approximately ∼1.5 cm of the ileum and colon were injected with 5 × 10^8^ *C. difficile* R20291 spores of strains wild-type (Δ*pyrE*/*pyrE*^+^) (SI *n* = 12; colon *n* = 10), Δ*bclA3* (SI *n* = 12; colon *n* = 12) and Δ*bclA3/bclA3*^+^ (SI *n* = 12; colon *n* = 11). **a**−**c** Representative confocal micrographs. *C. difficile* spores are shown in red, F-actin is shown in green and nuclei in blue (fluorophores colors were digitally reassigned for a better representation). The white arrows indicate internalized *C. difficile* spores, empty arrows indicate adhered *C. difficile* spore. Quantification of the spots (spores) number per 10^5^ µm^2^ relatives to wild-type of **d** adhered or **e** internalized in the ileum, and **f** adhered or **g** internalized in the colonic mucosa. **h** Schematics of the experimental design. Mice were infected with 5 × 10^7^ *C. difficile* spores strain R20291, wild-type (Δ*pyrE*/*pyrE*^+^) (*n* = 10), Δ*bclA3* (*n* = 16) or Δ*bclA3/bclA3*^+^ (*n* = 14) and were treated with vancomycin from day 3 to 7 and were monitored daily for **i**, relative weight during the R-CDI, **j** onset of diarrhea during the R-CDI. Spore adherence to the colonic tract was evaluated on day 11 to **k** medium colon. Error bars indicate the mean ± S.E.M. Statistical analysis was performed by **d**−**g** Mann-Whitney test; **j** Log-rank (Mantel-Cox) test ns indicates non-significant differences; * *P* < 0.05, \*\**P* < 0.001.

Next, to quantitatively assess the internalization of *C. difficile* spores into non-phagocytic cells, we developed an exclusion assay in which, in non-permeabilized cells, only extracellular spores are fluorescently labeled with anti-*C. difficile* spore antibody, while total spores can be quantified by phase-contrast microscopy; intracellular spores are not stained by anti-*C. difficile* spore antibody (absence of fluorescence) and are only detectable by phase-contrast microscopy (Extended Data Fig. 7a). With this assay, we probed that entry of 630 and R20291 spores into monolayers of Caco-2, T84, Vero, and HT29 cell lines significantly increased in the presence of FBS (Fig. 2i, j), as well as with serum from various mammalian species (Fig. 2k). The percentage of internalized spores of 630 and R20291 strains was highest at 5 h post-infection in Caco-2 and T84 cells (Extended Data Fig. 7b, c). Spores of various clinically relevant ribotypes were able to internalize into Caco-2 cells (Extended Data Fig. 7d). Overall, these results demonstrate that *C. difficile* spores are able to gain intracellular entry into non-phagocytic cells and that spore-entry is serum-dependent *in vitro*.

**Fig. 7.**
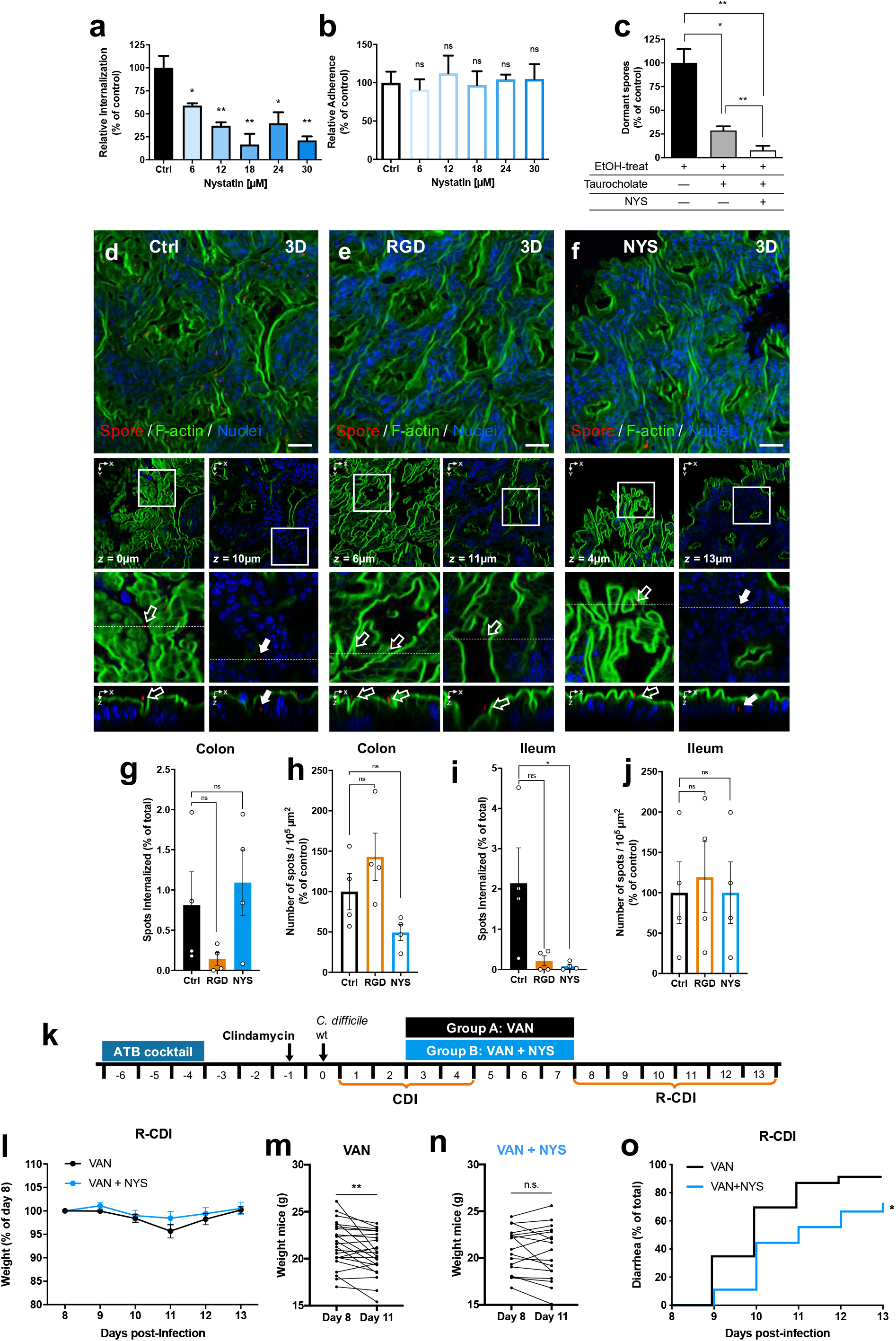
Nystatin reduces *C. difficile* spore internalization and reduces the R-CDI rates. **a** Internalization and **b** adherence of undifferentiated Caco-2 cells were pre-treated with 6, 12, 18, 24, and 30 μM of nystatin for 1 h and subsequently infected with *C. difficile* spores R20291 pre-incubated for 1 h with FBS. **c** Colony-forming units of spores on BHIS-CC with 0.1% of sodium taurocholate of a lysate of undifferentiated Caco-2 were pre-treated with 30 µM of nystatin (shown as NYS) or DMEM alone as control and infected with *C. difficile* spores, washed, and then treated with sodium taurocholate and ethanol (shown as EtOH). The cells were lysed and plated, so only spores that remain dormant after treating the cells with taurocholate (dormant) germinate in the plates. The number of CFU mL^−1^ was determined, and the percentage of adherence relative to the control. Loops of approximately ∼1.5 cm of the ileum and colon of C57BL/6 were injected with 3 × 10^8^ *C. difficile* R20291 with 250 nmol of RGD peptide (*n* = 4), or 17,000 UI kg^−1^ nystatin (*n* = 4) and saline (0.9% NaCl) as Ctrl (*n* = 4). **d**−**f** Representative confocal micrographs *C. difficile* spore is shown in red, F-actin is shown in green and nuclei in blue (fluorophores colors were digitally reassigned for a better representation). The white arrow indicates internalized *C. difficile* spores, empty arrow indicates adhered *C. difficile* spore. Quantification of spots number (spores) per 10^5^ µm^2^ of **g** internalized and **h** adhered spores in the ileum mucosa or **i** internalized and **j** adhered spores in the colonic mucosa in of C57BL/6. **k** Schematics of the experimental design of a mouse model of R-CDI. ATB cocktail treated C57BL/6 mice were infected with 6 × 10^7^ R20291 spores. The CDI symptoms were treated from days 3 to 7 with the inhibitor of spore internalization, nystatin and vancomycin (shown as VAN+NYS) (*n* = 18) or vancomycin alone as control (shown as VAN) (*n* = 23) and were monitored daily for **l** relative weight during the R-CDI. Weight loss comparison for animals treated with **m** vancomycin or **n** vancomycin and nystatin and **o** onset to diarrhea during the R-CDI. Error bars indicate the mean ± S.E.M. Statistical analysis was performed by **a**−**c**, Student’s *t*-test, **g**−**j**; Mann-Whitney test; **l**, Kruskal-Wallis, post-Dunn’s; **m, n**, Wilcoxon matched-pairs signed-rank test; **o**, Log-rank (Mantel-Cox) test; ns indicates non-significant differences, * *P* < 0.05, \*\**P* < 0.001.

### *C. difficile* spore-entry into intestinal epithelial cells requires Fn and Vn

Fn and Vn are extracellular matrix proteins, which are also present in mammal serum and are widely used by enteric pathogens to infect host cells^16,17^. We have shown previously that both, Fn and Vn, bind in a concentration-dependent manner to *C. difficile* spores^15^. To assess whether serum Fn and Vn contribute to *C. difficile* spore-entry, we evaluated the internalization assay in the presence of RGD peptide to block the interaction of Fn and Vn with their cognate receptors through the RGD binding domain^20,21^. RGD significantly reduced the extent of spore-entry into differentiated Caco-2 cells in the presence of human serum by ∼45% (Figure 3a, Extended Data Fig. 8a), indicating that serum Fn and Vn might be involved in *C. difficile* spore-entry in an RGD specific manner; by contrast, no decrease in adherence of *C. difficile* spores was evidenced in the presence of RGD (Fig. 3b, Extended Data Fig. 8b). Similar results were observed in undifferentiated Caco-2 cells (Extended Data Fig. 8c–f). We confirmed these results by showing that the infection with spores pre-incubated with Fn or Vn restored spore-entry into differentiated (Fig. 3c) and undifferentiated Caco-2 cells (Extended Data Fig. 9a), but had no impact in spore-adherence to differentiated (Fig. 3d) and undifferentiated Caco-2 cells (Extended Data Fig. 9b). Similar results were observed in differentiated Caco-2 cells pre-treated with Fn or Vn before infection with *C. difficile* spores (Extended Data Fig. 9c, d), confirming that the presence of Fn and Vn mediates *C. difficile* spore-entry.

### Intestinal barrier sites with accessible Fn and Vn

Both Fn and Vn are mainly located in the basal and basolateral membrane of epithelial cells where tight and adherent junctions are formed^16,17^. However, several the epithelial barrier suffers reorganization and/or disruption of tight and adherent junctions such as cell extrusion sites, goblet cells (GCs), at cell-cell junctions with neighboring cells and along villus epithelial folds^22-24^. Therefore, we hypothesized that sites undergoing adherent junction rearrangement also contained accessible Fn and Vn. We performed double staining, in which luminally accessible Fn and E-cadherin (Ecad), a marker for reorganization or disruption of adherent junctions^22^, were stained in non-permeabilized tissue. We first determined the relative number of IECs in the colonic tissue that expresses luminally accessible Ecad and found that nearly 16% of the IECs have this feature (Fig. 3h). Accessible Fn and Vn were observed in 27% and 14% of the IECs cells (Fig. 3i and 3k). We observed that most of the cells that had luminally accessible Fn or Vn also had accessible Ecad (Fig. 3j, l), and are likely undergoing major reorganization of the adherent junctions. However, a small fraction of epithelial cells with accessible Fn (33%) or Vn (12%) had no accessible Ecad (Fig. 3i, k). Luminally accessible Ecad has been previously found around mucus-expelling GCs in mice intestinal tissue^22^. Therefore, to quantify the relative abundance of GCs with accessible Ecad in our experimental conditions, we performed double immunostaining for accessible Ecad and the GC-specific marker Muc2^22,25^. We observed that 13% of the IECs were positive for Muc2 in colonic tissue (Fig. 3m) and all of Muc2-positive cells were positive for accessible Ecad (Fig. 3n). Luminally accessible Ecad has also been observed in mice ileal tissue^22^; we observed that nearly 17% of the IECs of mice ileal had luminal accessible Ecad (Extended data Fig. 10a, b). Next we quantified the relative abundance of GCs in the ileum mucosa and observed that nearly 9% of total IECs were positive for Muc2 (Extended Data Fig. 10c), of which 70% were positives for accessible Ecad (Extended Data Fig. 10d). This data supports the notion that Fn and Vn are also accessible in the intestinal epithelial barrier. Altogether, these results demonstrate the existence of sites in the intestinal barrier that undergo reorganization of adherent junctions that exhibit accessible Fn and Vn through which *C. difficile* spores can gain entry into the intestinal barrier.

### *C. difficile* spores internalize via Fn-α_5_β_1_ and Vn-α_v_β_1_ integrin *in vitro*

The Fn RGD loop between domains FnIII9 and FnIII10 enhances binding between Fn and α_5_β_1_ integrin^16^; Vn also has a similar RGD loop that enhances binding to α_v_β_1_ integrin^17^. To address whether binding of Fn and Vn to their cognate integrin receptors is required for *C. difficile* spore-entry into IECs, monolayers of Caco-2 cells were infected with *C. difficile* R20291 spores in the presence of the inhibitory RGD peptide, showing that in the presence of Fn or Vn, increasing concentrations of RGD progressively decreased spore-entry into differentiated (Fig. 4a, b) and into undifferentiated Caco-2 cells (Extended Data Fig. 11a, c), but not spore-adherence (Fig. 4c, d, Extended Data Fig. 10b, d) to Caco-2 cells. Next, through an antibody blocking assay, we assessed which integrin subunits are involved in Fn- and Vn-dependent entry of *C. difficile* spores into IECs. Results demonstrate that blocking the subunits of the collectin-binding, α_2_β_1_ integrin^26,27^ and β_3_ integrin subunit did not affect internalization nor adherence of *C. difficile* spores to Caco-2 cells in the presence of Fn (Fig. 4e, f) or Vn (Fig. 4g, h). However, a significant decrease in spore-entry, but not spore-adherence, to differentiated and undifferentiated Caco-2 cells was observed upon blocking each subunit of α_5_β_1_ integrin in the presence of Fn (Fig. 4e, f, Extended Data Fig. 11e, f), as well as blocking each subunit of α_v_β_1_ integrin in the presence of Vn (Fig. 4g, h, Extended Data Fig. 11g, h). These results were confirmed upon expressing each integrin subunit in Chinese hamster ovary (CHO) cells (Fig. 4i–k), a naïve cell line that otherwise does not express integrins. CHO cells expressing individual α_5_ or β_1_ integrin subunits exhibited significant spore-entry but not adherence in the presence of Fn (Fig. 4l, m); equally, CHO cells expressing individual α_v_ or β_1_ integrin subunit exhibited significant spore-entry but not spore-adherence in the presence of Vn (Fig. 4n, o). No increase in entry or adherence was detected in the absence of Fn and Vn (Extended Data Fig. 12 a, b). Altogether, these observations demonstrate that the internalization of *C. difficile* spores into IECs occurs through Fn-α_5_β_1_- and Vn-α_v_β_1_ uptake pathways.

### Fn and Vn bind to the hair-like extensions of *C. difficile* spores, formed by the collagen-like BclA3 exosporium protein

*C. difficile* spores of epidemically relevant strains exhibit hair-like projections that are likely to be formed by the collagen-like exosporium proteins^1,13^. Fn and Vn have a gelatin/collagen-binding domain^16,17^, suggesting that these molecules might interact with *C. difficile* spores through these hair-like projections. Indeed, through TEM coupled with immunogold labeling of Fn and Vn, we observed that more than ∼50% of the spores were positive for Fn- or Vn-immunogold particles (Extended Data Fig. 13a, b); immunogold Fn- and Vn-specific particles were observed in proximity to the hair-like extensions of *C. difficile* R20291 spores (Fig. 5a, b), suggesting that these structures might be implicated in spore-entry into IECs. Most epidemically relevant strains encode two collagen-like exosporium proteins, BclA2, and BclA3^1,13^. During the sporulation of R20291 strain, *bclA3* expression levels are ∼60-fold higher than those of *bclA2*^28^. Consequently, we first hypothesized whether BclA3 was responsible for the formation of the hair-like extensions. Therefore, we constructed a single *bclA3* mutant strain, in an epidemic R20291 background, by removing the entire gene through a *pyrE*-based allelic exchange system^29^ (Extended Data Fig. 14). Electron micrographs demonstrate that, as expected, wild-type R20291 (Δ*pyrE*/*pyrE*^+^) spores exhibited typical hair-like projections observed in previous reports^1,11,12^ (Fig. 5c). By contrast, the Δ*bclA3* deletion mutant formed spores that lacked the hair-like projections (Fig. 5d) that were restored upon complementation of the Δ*bclA3* mutant strain with a single wild-type copy of *bclA3* in the *pyrE* locus (Δ*bclA3*/*bclA3*^+^; Fig. 5e), indicating that BclA3 is required for the formation of these projections on the surface of *C. difficile* spores.

### BclA3 is required for Fn-α_5_β_1_- and Vn-α_v_β_1_-mediated spore-entry into IECs

To address whether BclA3 exosporium protein is implicated in *C. difficile* spore-entry into IECs, we first assayed whether the absence of BclA3 protein affected the internalization of *C. difficile* spores into IECs in the presence of Fn or Vn. As a control, we ensured that the anti-*C. difficile* spore goat serum used to quantify extracellular *C. difficile* spores, recognized Δ*bclA3* mutant spores (Extended Data Fig. 15a–c). Spores of the *C. difficile* Δ*bclA3* mutant strain exhibited a significant decrease in spore-entry into Caco-2 cells, but not adherence, to monolayers of Caco-2 cells was observed upon infection with *C. difficile* spores Δ*bclA3* mutant in the presence of Fn (Fig. 5f, g) and Vn (Fig. 5h, i). Importantly, the defect in spore-entry of the Δ*bclA3* mutant strain in the presence of Fn or Vn was restored to wild-type levels of internalization Δ*bclA3*/*bclA3*^+^ strain (Fig. 5f–i), indicating that BclA3 is required for Fn- and Vn-mediated internalization into IECs. We further confirmed these results in monolayers of HeLa cells, evidencing essentially identical results (Extended Data Fig. 16 a–d). Next, to address whether Fn-α_5_β_1_ and Vn-α_v_β_1_-mediated spore-entry is BclA3-specific, we carried out infection experiments with Δ*bclA3* mutant spores in monolayers of CHO cells expressing individual integrin subunits. In the presence of Fn, a significant decrease in spore-entry (Fig. 5j, k), but not in adherence (Extended Data Fig. 16e, f), was observed upon infection of CHO cells expressing the α_5_ or β_1_ integrin subunit with Δ*bclA3* mutant spores. Similarly, Δ*bclA3* mutant spores internalized to a significantly lesser extent than wild-type spores during infection of CHO cells expressing α_v_ or β_1_ integrin receptors in the presence of Vn (Fig. 5l, m); however, the absence of BclA3 had no impact on spore-adherence to CHO cells in the presence of Vn (Extended Data Fig. 16g, h). Fn- and Vn-mediated internalization of *C. difficile* spores into CHO cells expressing each integrin subunit was restored to wild-type levels upon infection with spores Δ*bclA3*/*bclA3*^+^ (Fig. 5j–m). Collectively, these results demonstrate that BclA3-Fn-α_5_β_1_- and BclA3-Vn- α_v_β_1_ are two pathways through which *C. difficile* spores can internalize into non-phagocytic cells.

### Inactivation of the exosporium protein BclA3 decreases spore-adherence, but not spore-entry, of *C. difficile* spores to the intestinal mucosa

To assess whether the collagen-like BclA3 exosporium protein also contributed to the internalization of *C. difficile* spores into the intestinal mucosa *in vivo*, we used a colonic and ileal loop mouse model (Fig. 6a-c, Extended data Fig. 17 a-c). In contrast to our *in vitro* data, analysis of colonic mucosa sections show that inactivation of *bclA3* leads to a significant decrease of ∼60% in spore-adherence per 10^5^ µm^2^ to the ileum mucosa (Fig. 6d); however, no differences were observed in spore internalization relative to the total adhered-spores (Fig. 6e). A similar trend was evidenced in ileal loops, where Δ*bclA3* spores adhered in a ∼50% lower than wild-type spores per 10^5^ µm^2^ (Fig. 6f); however, no differences were observed in spore internalization relative to total adhered-spores (Fig. 6g). The defects in spore-adherence to the colonic and ileum mucosa were restored to wild-type levels upon complementing the wild-type *bclA3* allele in the Δ*bclA3* mutant (Fig. 6d–g, Extended data Fig. 17 a-c). Strikingly, these data indicate that the collagen-like BclA3 exosporium protein is required for *C. difficile* spore adherence to the intestinal mucosa, and that additional spore-surface proteins are contributing to redundant spore-entry pathways *in vivo*.

### The *C. difficile* collagen-like BclA3 exosporium protein contributes to spore-persistence and recurrence of the disease in mice

Since BclA3 is essential for spore-entry into IECs *in vitro* and for adherence in the intestinal mucosa *in vivo*, we hypothesized that BclA3 might mediate spore-persistence and contribute to R-CDI. Therefore, antibiotic-treated mice were infected with spores of wild-type (Δ*pyrE*/*pyrE*^+^), Δ*bclA3* and Δ*bclA3*/*bclA3*^+^ strains (Fig. 6h). All three groups of mice exhibited similar weight loss during the initiation of CDI, and all manifested signs of diarrhea within 3 days post-infection (Extended Data Fig. 17d, e). Similar levels of *C. difficile* spores shed in feces were observed during the initiation of CDI between mice infected with wild-type (Δ*pyrE*/*pyrE*^+^), and the mutants Δ*bclA3* and Δ*bclA3*/*bclA3*^+^ (Extended Data Fig. 17f). These results indicate that the absence of BclA3 does not affect the initiation of CDI. Next, the impact of BclA3 in the recurrence of the infection was assessed by treating *C. difficile* infected mice with vancomycin for 5 days (Fig. 6h), R-CDI was monitored from day 8 post-infection. No significant differences in weight loss were evidenced after vancomycin treatment (days 8-11; Fig. 6i). However, a significant delay in the onset of diarrhea during R-CDI was observed after vancomycin treatment in mice infected with Δ*bclA3* mutant spores compared to wild-type infected-mice (Fig. 6j). The defect in R-CDI observed in Δ*bclA3* mutant infected-mice was restored to wild-type levels with the complemented Δ*bclA3/bclA3*+ strain (Fig. 6j). Although there were no significant differences in the levels of *C. difficile* spores, shed in the feces during R-CDI (Extended Data Fig. 17f), significantly lower CFUs of Δ*bclA3* mutant spores were detected in the medium colon compared to wild-type spores (Fig. 6k) but not in other sections of the intestinal tract (Extended Data Fig. 17g–i). Again, this defect was reverted in the complemented Δ*bclA3/bclA3*+ strain (Fig. 6k). Cytotoxicity levels in the cecum content of Δ*bclA3* mutant-infected mice were similar to those found in animals infected with wild-type strain (Extended Data Fig. 17j). Collectively, these results demonstrate that the collagen-like BclA3 exosporium protein is involved in *C. difficile* persistence and recurrence of the disease.

### Inhibition of spore-entry into IECs renders *C. difficile* spores susceptible to taurocholate-germination

Since *C. difficile* spore-entry into IECs requires integrin receptors, we tested whether cholesterol-lipid rafts, commonly required by integrin receptors for endocytosis^30,31^, were also required for uptake of *C. difficile* by IECs. Therefore, we used the cholesterol-chelating agent, nystatin, which is a caveolin-related pathway inhibitor that disrupts membrane microdomains known to be implicated in integrin-mediated endocytosis and pathogen uptake^30,32^. Cells were preincubated with nystatin for 1 h at 37 °C and infected in the same medium containing the inhibitor and *C. difficile* spores. *C. difficile* spore-entry was inhibited in a dose-dependent manner into Caco-2 cells and T84 cells (Fig 7a, b, Extended Data Fig. 18a, b). 30µM of nystatin inhibited the spore-entry by about 80% in human cell lines Caco-2 and about 65% in T84 cells. We determined cell viability in the presence of nystatin by MTT at the highest concentration of the inhibitor. Cell viability was generally around 90% (Extended Data Fig. 18c). These results suggest that *C. difficile* spore-entry is sensitive to cholesterol-sequestering compounds.

Vancomycin administration leads to increased fecal-concentration of primary bile acids^33^ leading to enhanced *C. difficile* spore germination^34,35^, suggesting that luminal taurocholate would trigger germination of extracellular *C. difficile* spores that could subsequent become inactivated by vancomycin. Therefore, we hypothesize that intracellular spores should remain dormant in the presence of taurocholate. To test this hypothesis, monolayers of 1-h nystatin-treated or untreated Caco-2 cells were infected for 3h with serum-treated *C. difficile* spores. Next, infected monolayers were washed and treated with taurocholate to trigger germination of extracellular spores, followed by ethanol-treatment to inactivate germinated *C. difficile* spores. We observed that not all of the spores became ethanol-sensitive upon taurocholate-treatment of infected Caco-2 monolayers (Fig. 7c), suggesting that internalized spores were protected from taurocholate-triggered germination. We confirmed this by evidencing a significant increase in ethanol-sensitive germinated spores in the presence of nystatin (Fig. 7c). These results indicate that blocking *C. difficile* spore-entry contributes to taurocholate-triggered germination of *C. difficile* spores and subsequent spore-inactivation.

### Inhibition of *C. difficile* spore-entry into the intestinal barrier reduces recurrence of the disease in mice

To address whether *in vivo C. difficile* spore-entry into the intestinal barrier also required RGD-binding integrins^36^, colonic and ileal loop assays were assessed in the presence of RGD during *C. difficile* spore infection (Fig. 7d, e, Extended data Fig 18d, e). Ileal and colonic loops were injected with RGD peptide and *C. difficile* spores during 5h, then were processed and visualized in confocal microscopy. Consistent with our *in vitro* data, in the colonic loop sections, we observed that presence of RGD peptide reduced spore internalization by ∼82% (Fig. 7g), while no difference in spore-adherence was observed (Fig. 7h); similarly, in ileal loop sections, we observed that RGD peptide decreased spore internalization by ∼90% (Fig. 7i) and does not affect the spore adherence to the ileum mucosa (Fig. 7j). These results demonstrate that *C. difficile* spore-entry *in vivo* is RGD-binding integrin-dependent.

Since the RGD-dependency of spore-entry into the intestinal barrier is likely attributed to integrin receptors, we address whether the cholesterol-sequestering drug, nystatin, could block internalization of *C. difficile* spores into the intestinal barrier *in vivo* in the colonic and ileum mouse mucosa. Mice were treated for 24-h with nystatin or saline as a control prior to surgery and during intestinal loop-infection (Fig. 7d, f, Extended data Fig 18d, f) then were infected with *C. difficile* spores for 5h, then tissues were processed for confocal microscopy. In the colonic loop section, we observed that nystatin had no effect on spore internalization (Fig. 7g) and in spore-adherence to the colonic mucosa (Fig. 7h); however, in the ileal loop sections, the presence of nystatin significantly decreased spore internalization by ∼96% (Fig. 7i), and no effect in *C. difficile* spore-adherence to the ileum mucosa was observed (Fig. 7j).

Since *C. difficile* spore-entry prevents taurocholate-germination, contributing to the persistence of *C. difficile* spores during the disease; we hypothesized that administration of the inhibitor of spore-entry, nystatin, during CDI-treatment with vancomycin, could reduce the recurrence of the infection in a previously developed mouse model of R-CDI^8^ (Fig. 7k). To address this question, antibiotic-treated mice were infected with *C. difficile* R20291 spores. During the first episode of CDI, both groups of mice had similar levels of weight loss, the timing of the onset of diarrhea and shed similar amounts of *C. difficile* spores during the initiation of CDI (days 1-3) (Extended Data Fig.18g–i). At day 3 post-infection, animals were treated with vancomycin or a mixture of vancomycin and nystatin for 5 days (Fig. 7k). Vancomycin-treated mice exhibited a significant decrease in weight during R-CDI, which became highest at day 11 post-infection (4^th^ day after vancomycin treatment; Fig. 7k, m). By contrast, CDI-animals treated with vancomycin and nystatin had no significant decrease in weight loss during the recurrence of the infection (Fig. 7k, n). These observations were confirmed upon monitoring the onset of diarrhea during R-CDI (Fig. 7o), where we observed a significant delay in the onset of recurrent diarrhea in CDI-mice treated with the mixture of vancomycin and nystatin compared to vancomycin alone (Fig. 7o). The animals shed similar amounts of *C. difficile* spores during R-CDI (Extended Data Fig. 18i). Collectively, these results demonstrate that the administration of a pharmacological inhibitor of internalization of *C. difficile* spores during vancomycin treatment delays the incidence of recurrence of the infection.

## Discussion

During CDI, *C. difficile* spore-formation is essential in the recurrence of the disease^7^, yet the underlying mechanisms that correlate *C. difficile* spore-persistence and recurrence of the disease remain unclear. In this study, we unravel a novel and unexpected mechanism employed by *C. difficile* spores to interact with the intestinal mucosa that contributes to the recurrence of disease. Our results have identified host molecules, cellular receptors, and a spore-surface ligand involved in spore-entry into IECs. Importantly, intracellular spores remain dormant in the presence of germinant. Using nystatin, a pharmacological inhibitor of spore-entry in combination with antibiotic treatment, leads to a reduction in the recurrence of the disease in mice. Together, these observations open a new angle for therapeutic interventions of CDI to prevent the recurrence of the disease.

Our results identified host molecules and cellular receptors involved in the entry of *C. difficile* spores into IECs. The presence of Fn or Vn allows *C. difficile* spores to gain intracellular access to IECs, in an RGD-specific manner, and through specific integrin receptors (i.e., α_5_β_1_ and α_v_β_1_). These observations were confirmed by the *in vivo* inhibition of *C. difficile* spore-entry in the presence of the RGD peptide, which specifically inhibits interactions between Fn-α_5_β_1_ and Vn-α_v_β1^20,21^. Although Fn and Vn are mainly located in the basal and basolateral membrane of IECs contributing to cell polarity^16,17^, antibody staining of healthy ileum and colonic tissue demonstrate that Fn and Vn are luminally accessible in a significant fraction of the IECs. Most of these cells were positive for luminally accessible Ecad, and suggests that these cell types include cell-extrusion, cells next to extrusion sites, and epithelial folds that typically undergo adherent junction reorganization^22-24^. However, a small fraction of cells positive for luminally accessible Fn and Vn were negative for luminally accessible Ecad, suggesting a novel phenotype within cells at the intestinal epithelial barrier. We also confirmed previous observations in mice that identified GCs have luminally accessible Ecad^22^, suggesting that these cell types might also be targeted by *C. difficile* spores to gain entry into the epithelial barrier. M cells are an additional cell-type that might contribute to *C. difficile* spore-entry into the intestinal epithelial barrier includes since they expresses β_1_-integrin at the apical surface in contrast to its normal basolateral location in enterocytes^22,37,38^. The fact that *in vivo* spore entry was RGD-binding integrin-specific, suggests that Fn and Vn are accessible and employed by *C. difficile* spores to gain entry into IECs, which is consistent with the presence of accessible Fn and Vn in ileal and colonic loops. It is noteworthy that while RGD-specific entry was observed in both ileal and colonic loops, nystatin was only able to reduce spore-entry into the ileum, but not colonic mucosa. This suggests that caveolae-independent endocytosis of *C. difficile* spores might prime in the colonic epithelia. During CDI, C. *difficile* toxins disrupt adherent junctions, leading to progressive exposure of deep regions of the colonic epithelium as infection advances. One consequence of this cellular disorganization may be an alteration to the distribution of cell receptors that may lead to increased adherence and internalization of *C. difficile* spores into the intestinal mucosa. Together, these observations prompt further studies to address how epithelium remodeling contributes to persistence of *C. difficile* spores and recurrence of the disease.

Another major contribution of this work is the role of the spore surface collagen-like BclA3 exosporium protein in *C. difficile* spore-entry into IECs in a Fn-α_5_β_1_- and Vn-α_v_β_1_- dependent manner. Our previous work shows that Fn and Vn bind in a dose-dependent manner to *C. difficile* spores^15^. By immunogold-electron microscopy, our results demonstrate that Fn and Vn bind to the hair-like projections of *C. difficile* spores. We also demonstrate that they are formed by the collagen-like exosporium glycoprotein BclA3. It is noteworthy that experiments with monolayers of Caco-2 cells and CHO cells expressing integrin subunits demonstrate that BclA3 is essential for spore-entry in the presence of Fn and Vn in a integrin-dependent manner; results that contrast with BclA3 being essential for adherence to the intestinal mucosa, but not for spore-entry into the intestinal barrier. Coupling these results with those of *in vivo* RGD-specific *C. difficile* spore-entry into the intestinal barrier, indicates that additional spore-surface proteins might play redundant roles during *in vivo* spore-entry. Regardless of these incongruencies, we observed that BclA3 contributes to the recurrence of the disease in a mouse model, suggesting that BclA3-mediated spore-adherence to the intestinal mucosa might contribute to spore persistence and recurrence of the disease. The differences in spore-adherence to the colonic tissue after R-CDI observed in the medium colonic tissue of mice might relate to the absence of mucosal folds typically observed in the distal and proximal colon of mice^39^. Here, we have shown that BclA3 uses Fn and Vn and their specific integrins to gain-entry into IECs and that BclA3 is essential for *C. difficile* spore adherence to the intestinal mucosa and contributes to the recurrence of the disease.

The work presented here also shows that *C. difficile* spore-entry into IECs contributes to spore dormancy in the presence of primary bile salts (i.e., taurocholate), and that blocking *in vivo* spore-entry during antibiotic treatment (vancomycin), leads to reduced recurrence of CDI in mice. This brings a broader understanding of how strict anaerobic spore-formers can persist in the host and remain dormant in a dysbiosis environment enriched with bile acids that trigger spore germination. Intracellular bacterial spores may survive until released back to the luminal environment to recolonize the host. Although the precise mechanism of how intracellular spores would contribute to the recurrence of the disease is unclear and prompts further studies, it may involve the rapid renewal of the intestinal epithelium, which, due to rapid proliferation and differentiation of multipotential stem cells located in the crypts of Lieberkühn^40-43^, renew the epithelial barrier every 5 days. The factors that contribute to infection recurrence, although partly linked to continued disruption of the microbiota^44^, are also directly linked to the persistence of *C. difficile* spores in the host. This is particularly relevant for CDI, considering that the rates for recurrent CDI are around ∼18 to 32%, and may rise between 45 and 65% during subsequent recurrent episodes^2,44^. Importantly, this *C. difficile* spore-entry phenotype provides an additional point of intervention of disease recurrence and therapeutic susceptibility. The cholesterol-sequestering drug, nystatin, is FDA approved for oral administration^45^, raising new approaches to develop pharmacological formulations that target *C. difficile* spore-entry during disease. Similarly, BclA3 and α_5_β_1_ and α_v_β_1_ integrins are also candidates drug targets to combat recurrent *C. difficile* infections.

## Methods

### Data reporting

No statistical methods were used to predetermine the sample size. The experiments were not randomized, and investigators were not blinded to allocation during experiments and outcome assessment.

### Bacterial strains and growth conditions

*C. difficile* strains (See Table S1) were routinely grown at 37 °C under anaerobic conditions in a Bactron III-2 anaerobic chamber (Shellab, USA) in BHIS medium: 3.7% weight vol^−1^ brain heart infusion broth (BD, USA) supplemented with 0.5% weight vol^−1^ yeast extract (BD, USA) and 0.1% weight vol^−1^ L-cysteine (Merck, USA) or on BHIS agar plates. *E. coli* strains were routinely grown aerobically at 37 °C under aerobic conditions with shaking (200 r.p.m.) in Luria-Bertani (LB) medium (BD, USA), supplemented with 25 µg mL^−1^ chloramphenicol (Merck, USA), where appropriate.

For mutant construction, a defined *C. difficile* minimal medium (CDMM) media was prepared as described by Cartman and Minton^46^ as an uracil-free medium when performing genetic selections. For CDMM broth preparation, 5× amino acids (50 mg mL^−1^ casamino acids, 2.5 mg mL^−1^ L-tryptophan, 2.5 mg mL^−1^ L-cysteine), 10× salts (50 mg mL^−1^ Na_2_HPO_4,_ 50 mg mL^−1^ NaHCO_3_, 9 mg mL^−1^ KH_2_PO_4_, 9 mg mL^−1^ NaCl), 20× glucose (200 mg mL^−1^ D-glucose), 50× trace salts (2.0 mg mL^−1^ (NH_4_)_2_SO_4_, 1.3 mg mL^−1^ CaCl_2_·2H_2_O, 1.0 mg mL^−1^ MgCl_2_·6H_2_O, 0.5 mg mL^−1^ MnCl_2_·4H_2_O, 0.05 mg mL^−1^ CoCl_2_·6H_2_O), 100× iron (0.4 mg mL^−1^ FeSO_4_·7H_2_O) and 100× vitamins (0.1 mg mL^−1^ D-biotin 0.1, mg mL^−1^ calcium-D-pantothenate, 0.1 mg mL^−1^ pyridoxine) stock solutions were made by dissolving their components in Milli-Q water and filter sterilizing (0.2-μm pore size) prior to use. Solutions were mixed to obtain a final CDMM media made of 10 mg mL^−1^ casamino acids, 0.5 mg mL^−1^ L-tryptophan, 0.5 mg mL^−1^ L-cysteine, 5 mg mL^−1^ Na_2_HPO_4,_ 5 mg mL^−1^ NaHCO_3_, 0.9 mg mL^−1^ KH_2_PO_4_, 0.9 mg mL^−1^ NaCl, 10 mg mL^−1^ D-glucose, 0.04 mg mL^−1^ (NH_4_)_2_SO_4_, 0.026 mg mL^−1^ CaCl_2_·2H_2_O, 0.02 mg mL^−1^ MgCl_2_·6H_2_O, 0.01 mg mL^−1^ MnCl_2_·4H_2_O, 0.001 mg mL^−1^ CoCl_2_·6H_2_O, 0.004 mg mL^−1^ FeSO_4_·7H_2_O, 0.001 mg mL^−1^ D-biotin, 0.001 mg mL^−1^ calcium-D-pantothenate and 0.001 mg mL^−1^ pyridoxine^46^. For solid medium, agar (BD, USA) were mixed with CDMM to a final concentration of 1.0% weight vol^−1^. Finally, media were supplemented with uracil (Sigma–Aldrich, USA) at 5 mg mL^−1^ and 5-Fluoroorotic acid (5-FOA) (USBiological USA) at 2 mg mL^−1^ as described^47,48^.

### Cell Lines and Reagents

Caco-2, Vero, HT29, and Chinese hamster ovary (CHO) were obtained from ATCC (USA). Dr. Mauricio Farfán (Universidad de Chile, Chile) gently provided T84 cells. Caco-2 and Vero were routinely grown at 37 °C with 5% of CO_2_ with Dulbecco’s modified Eagle’s minimal essential medium (DMEM) High Glucose (HyClone, USA); CHO cells in Ham’s F-12K (Kaighn’s) medium; T84 in DMEM/F12 1:1 (HyClone, USA); and HT29 in RPMI 1640. All media were supplemented with 10% vol vol^−1^ inactivated Fetal Bovine Serum (FBS) (HyClone, USA) and 100 U ml^−1^ penicillin, and 100 µg ml^−1^ streptomycin (HyClone, USA). T84 cells were cultured onto Transwell (Corning USA) until 1,000**–**2,000 Ω. For transfected CHO cells (CHO-α_v_, CHO-α_5,_ and, CHO-β_1_), the culture media was supplemented with 1,500 µg mL^−1^ geneticin (HyClone USA). For immunofluorescence experiments, cells were plated over glass coverslip in a 24-wells plate and cultured for 2-days post confluence (undifferentiated) or 8-days post-confluence (differentiated), changing the culture medium every other day.

### Spore Preparation

Spores preparation was done as previously has been published^18^. Briefly, 100 μL of 1:1,000 dilution of an overnight culture in BHIS was plated in 70:30 agar plates that were prepared as follow: 6.3% weight vol^−1^ (BD, USA), 0.35% weight vol^−1^ protease peptone (BD, USA), 0.07% ammonium sulfate (NH_4_)_2_SO_4_ (Merck USA), 0.106% weight vol^−1^ Tris base (Omnipur, Germany), 1.11% weight vol^−1^ brain heart infusion extract (BD, USA) and 0.15% weight vol^−1^ yeast extract (BD, USA), 1.5% weight vol^−1^ Bacto agar (BD, USA). Plates were incubated for 7 days at 37 °C under anaerobic conditions in anaerobic chamber Bactron III-2 (Shellab USA). Then plates were removed from the chamber, and colonies were scraped out with ice-cold sterile Milli-Q water. Then the sporulated culture was washed five times with ice-cold Milli-Q water in micro-centrifuge at 18,400×*g* for 5 min each. To separate spores, the sporulated culture was loaded in 45% weight vol^−1^ autoclaved Nycodenz (Axell USA) solution and centrifugated at 18,400×*g* for 40 min. Spore pellet was separated and washed 5 times at 18,400×*g* for 5 min with ice-cold sterile Milli-Q water to remove Nycodenz. Spores were counted in Neubauer chamber, and volume adjusts at 5 × 10^9^ spores mL^−1^ and stored at −80 °C.

### *C. difficile* mutant construction by allelic exchange

Primer design and amplification of *C. difficile* R20291 strain were based on the available *C. difficile* genomes from the EMBL/GenBank databases with accession number FN545816. The oligonucleotides and the plasmids/strains used in this study are listed in Table S1 and Table S2, respectively. In-frame deletions in *C. difficile* R20291 were made by allelic exchange using *pyrE* alleles^47^.

To remove the *bclA3* gene, a 1086 bp allelic exchange cassette was obtained by overlap extension PCR of the LHA and RHA originated by amplification with primer pairs P332 (FP-LHA-bclA3-pyrE)/P334 (RP-LHA-bclA3-pyrE) and P335 (FP-RHA-bclA3-pyrE)/P336 (RP-RHA-bclA3-pyrE), each of 544 bp and 542 bp in size. The resulting cassette yielded complete removal of the entire *bclA3* cassette. Next, this cassette was cloned into Sbf1/AscI sites in pMTL-YN4, giving plasmid pDP376. To verify the correct construction of the plasmids, all constructs were Sanger sequenced.

The plasmids obtained were transformed into *E. coli* CA434 (RP4) and mated with *C. difficile* R20291 Δ*pyrE*^47^. *C. difficile* transconjugants were selected by sub-culturing on BHIS agar containing 15 µg mL^−1^ thiamphenicol (Sigma–Aldrich USA) and 25 µg mL^−1^ cefoxitin (Sigma–Aldrich USA) and re-streaked five times. The single-crossover mutants identified were streaked onto *C. difficile* minimal medium (CDMM) ^49^ with 1.5% weight vol^−1^ agar supplemented with 2 mg mL^−1^ 5-Fluoroorotic acid (USBiological, USA) and 5 µg mL^−1^ uracil (Sigma–Aldrich USA) in order to select for plasmid excision. Confirmation of plasmid excision was made by negative selection in BHIS-thiamphenicol plates. The isolated FOA-resistant and thiamphenicol resistant colonies were screened using the primer pair P664 (FP-bclA3-detect) / P665 (RP-bclA3-detect) for the *bclA3* mutant. All mutants were whole-genome sequenced to confirm the genetic background and that no additional SNPs were introduced during the genetic manipulation. For correction of the *pyrE* mutation, transconjugants with pMTL-YN2C were streaked onto minimal media without uracil or FOA supplementation, and developed colonies were analyzed further.

### Complementation by allelic exchange at the *pyrE* locus

To complement the Δ*bclA3* mutation, a 3,564 bp fragment containing 372 bp upstream of the start codon of *bclA3* and the entire bicistronic operon formed by *sgtA* and *bclA3* was PCR amplified with primer pairs P476 (NFP-bclA3c-promotor)/P477 (NRP-bclA3c) and cloned into BamHI/EcoRI sites of pMTL-YN2C, giving plasmid pMPG1. Next, plasmid pMPG1 was transformed into *E. coli* CA434 and subsequently conjugated with *C. difficile* R20291Δ*pyrE* Δ*bclA3*, respectively, as described above. The transconjugants obtained were streaked onto CDMM and tested by colony PCR using primer pair P530 (Fp-*pyrE* detect)/P529 (RP-*pyrE* detect) for *pyrE* reversion. Complemented strains were also subjected for whole-genome sequencing.

### Transfections of CHO cells with α5, αv and, β1 integrins

The integrins subunits were overexpressed in CHO cells line with the following plasmids: Alpha 5 integrin-GFP (Addgene plasmid# 15238)^50^, miniSOG-Alpha-V-Integrin-25 (Addgene plasmid # 57763)^51^ and Beta1-GFP in pHcgreen donated by Martin Humphries (Addgene plasmid # 69804)^52^. CHO cells were seeded on coverslips in 24-well plates until reach 70-90% of confluency and were transfected using Lipofectamine® LTX (Invitrogen USA) according to manufacturer protocol with 1 µg of each plasmid. Transfected cells were analyzed and confirmed by positive GFP fluorescence in epifluorescence microscopy. When the population GFP positive cells were higher than 50%, were selected for geneticin resistance with 1,500 µg mL^−1^ of geneticin until ∼100% of GFP positive cells, and the level of expression of the integrin subunits in the cells was confirmed by Western blot.

### SDS-PAGE and Western blot of transfected CHO cells

Transfected CHO cells were washed and homogenized with RIPA buffer that was prepared as follow: 50 mM buffer Tris HCl (Omnipur, Germany); 150 mM NaCl (Sigma–Aldrich, USA); 0.5% weight vol^−1^ deoxycholate (Sigma–Aldrich, USA); 1% vol vol^−1^ NP 40 (Sigma–Aldrich, USA); 1 mM EGTA (Sigma–Aldrich, USA); 1 mM EDTA (Sigma–Aldrich); 0.1% weight vol^−1^ SDS (Winkler, USA). The cell lysate was centrifuged at 18,400×*g* for 30 min at 4 °C, and protein concentration was quantified by BCA protein kit (RayBiotech USA). Next, 20 µg of protein were suspended in 2X SDS-PAGE sample loading buffer, boiled and electrophoresed on 12% vol vol^−1^ and 4% vol vol^−1^ acrylamide SDS-PAGE gels (Bio-Rad Laboratories, Canada) on MiniProtean® camera (Bio-Rad Laboratories, Canada). Then proteins were transferred to a nitrocellulose membrane (Bio-Rad Laboratories, Canada). Membranes were blocked then probed in Tris-Buffered saline containing 0.1% vol vol^−1^ Tween (TTBS), with 2% weight vol^−1^ BSA, incubated with mouse anti-α_v_, α_5_ and β_1_ antibody (SC166665, SC376156 y SC374429; Santa Cruz Biotechnologies, USA) and 1:1,000 mouse anti-alpha tubulin (T5168 Sigma–Aldrich USA) in 2% BSA-TTBS as loading control at concentration and were washed 3 times with TTBS. Membranes were incubated with 1:10,000 vol vol^−1^ secondary antibody anti-mouse horseradish peroxidase (HRP) conjugate (A5278, Sigma–Aldrich, USA) in 2% weight vol^−1^ BSA-TTBS. HRP activity was detected with a chemiluminescence detection system (Fotodyne Imaging system, USA.) by using PicoMax sensitive chemiluminescence HRP substrate (Rockland Immunochemicals, USA.).

### Germination assay of extracellular *C. difficile* spores in Caco-2 cells

Two–day old confluent monolayers of Caco-2 cells were treated with 30 µM of nystatin for 1 h at 37 °C or DMEM high glucose without FBS as control. To infect cells, *C. difficile* spores at an MOI of 10 were pre-incubated 1 h at 37 °C with 20 µL of NHS and then suspended in 200 µL that were added to each well; FBS final concentration 10% vol vol^−1^

To remove unbound spores, monolayers were washed 3 times with PBS, and cells were incubated with 0.1% weight vol^−1^ sodium taurocholate (Sigma–Aldrich, USA) in DMEM for 1 h at 37 °C (or DMEM as control) and washed 3 times with PBS. Cells were treated with 100% ethanol for 10 min, and cells were lysed with PBS-0.06% Triton X-100 for 10 min, plated in BHIS-CC supplemented with 0.1% weight vol^−1^ sodium taurocholate and incubated at 37 °C overnight. The number of CFU mL^−1^ was determined, and the percentage of adherence relative to the control.

### Infection of monolayers of cell lines with *C. difficile* spores

Adherence and intracellular spores were examined using a differential immunofluorescence staining procedure, as previously described^14,53^ with modifications. To evaluate the dynamic of *C. difficile* spore in Caco-2 cells and T84 cells. Caco-2 and T84 cells were grown on coverslips in 24–wells tissue culture plates until they reach a monolayer of 2-days post-confluency and were infected for 0.5, 1, 3, 5, and 8 h at 37 °C at an MOI of 10 of *C. difficile* spores pre-incubated 1h at 37 °C with 20µL of FBS and then suspended in the infection volume of 200µL that was added to each well. Then were washed gently in PBS prior to immunostaining as described below.

To evaluate if *C. difficile* internalize in different cell lines, undifferentiated, differentiated Caco-2, T84, Vero and HT29 cells were infected at 37 °C with an MOI 10 with *C. difficile* spore strain 630, and R20291 preincubated 1h at 37 °C with FBS or DMEM as control as was described above. Were washed gently in PBS prior to immunostaining, as described below.

Also, undifferentiated Caco-2 cells were infected at an MOI of 10 with *C. difficile* spores preincubated 1h at 37 °C with FBS, mouse serum (Pacific Immunology, USA), rat serum (Pacific Immunology, USA), rabbit serum (Pacific Immunology, USA) and NHS (Complement Technology USA) as was described above. Then were washed gently in PBS prior to immunostaining, as described below.

To evaluate if *C. difficile* internalization occurs in different strains, undifferentiated Caco-2 cells were infected at an MOI of 10 with *C. difficile* spores R20291, M120 and spores of *C. difficile* clinical isolates PUC52, PUC30, PUC 25, PUC31, PUC 98 and PUC 131^54^, which were pre-incubated 1h at 37 °C with FBS as was described above. Then were washed gently in PBS prior to immunostaining, as described below.

To assess that the internalization of *C. difficile* spores into IECs is through the specific interaction between Fn or Vn and their cognate integrin receptors, infection experiments were done in the presence of the RGD peptide ^20,21^. Briefly, differentiated and undifferentiated Caco-2 cells were incubated with 0, 1, 3, and 5 µg mL^−1^ of RGD peptide (Abcam USA) for 1 h, 37 °C then were infected for 3h at 37 °C with spores pre-incubated 1h at 37 °C with NHS as was described above, then samples were washed gently in PBS prior to immunostaining as described below.

To evaluate whether the internalization of the spores is mediated by Fn and Vn, differentiated and undifferentiated Caco-2 cells were treated for 1h 37 °C at with 10 µg mL^−1^ of purified human Fn or human Vn in DMEM and then were infected with an MOI 10 of untreated *C. difficile* spores R20291. Also, the infection was performed using untreated Caco-2 cells that were infected for 3h at 37 °C with an MOI 10 of *C. difficile* spores pre-incubated for 1h at 37 °C with 10 µg mL^−1^ of human Fn or human Vn in DMEM. Then samples were washed in PBS prior to the immunostaining as described below.

To confirm that the internalization of *C. difficile* spores is dependent of Fn and Vn, we perform an infection assay in differentiated and undifferentiated Caco-2 cells that were pre-incubated with 1, 3, or 5 µg mL^−1^ of RGD peptide and were infected with an MOI of 10 with *C. difficile* spores pre-incubated for 1h 37 °C with 10 µg mL^−1^ of purified human Fn or human Vn in DMEM. Then samples were washed in PBS prior to the immunostaining as described below.

To identify the integrin subunits implicated in spore-entry, an antibody blocking assay was performed using mouse monoclonal antibodies against individual integrin subunits: anti-human integrin α_5_, α_v_, (ab78614, ab16821; Abcam, USA), α_2_, β_1_ and β_3_ (MAB1950Z, MAB1959Z, and MAB2023Z; Millipore USA); and control non-immune IgG antibody (I5006, Sigma–Aldrich, USA). Caco-2 cells were incubated with 200µL of DMEM with the appropriate antibodies at 5 µg mL^−1^ for 1 h at 37 °C. The cells were infected for 3h at 37 °C with spores pre-incubated for 1h at 37 °C with 10 µg mL^−1^ of purified human Fn or human Vn in DMEM. Then samples were washed in PBS prior to the immunostaining as described below.

In order to demonstrate that *C. difficile* spore entry requires the integrins subunits α_5_, α_v,_ and β_1,_ Then CHO cells with ectopic expression of α_5_, α_v_ and, β_1_ integrins, were infected 3 h at 37 °C with an MOI of 10 of *C. difficile* spores that were preincubated with 10 µg mL^−1^ of purified human Fn or human Vn in DMEM. Then samples were washed in PBS prior to the immunostaining as described below.

To evaluate whether the collagen-like exosporium protein BclA3 is required for *C. difficile* spore entry into intestinal epithelial cells is dependent of differentiated Caco-2 cells and monolayers of HeLa cells, were infected for 3 h at 37 °C with an MOI of 10 with wild-type (Δ*pyrE*/*pyrE*^+^), Δ*bclA3* and Δ*bclA3/bclA3+ C. difficile* R20291 spores that were preincubated for 1h at 37 °C with 10 µg mL^−1^ of purified human Fn or human Vn in DMEM. Then samples were washed in PBS prior to the immunostaining as described below.

To evaluate the effect of nystatin in the internalization of *C. difficile* spores, Caco-2 cells were pre-incubated with 6, 12, 18, 24, and 30 μM nystatin or T84 cells were incubated with 30µM of nystatin (Sigma–Aldrich USA) for 1 h at 37 °C in DMEM and in the same media were infected for 3 h at 37 °C with spores at an MOI 10 pre-incubated with FBS as was described above. At the used concentration of nystatin, the cell viability of treated Caco-2 cells and T84 for 4 h was ∼90%, as was observed by trypan blue (Invitrogen USA) and MTT assay (Life Technologies USA) according to manufacturer protocols.

### Immunofluorescence of adhered *C. difficile* spores in infected monolayers and epifluorescence analysis

The aforementioned infected monolayers of cells were subsequently fixed with PBS–4% paraformaldehyde for 10 min, then were washed 3 times and blocked with PBS–1% BSA overnight at 4 °C; in nonpermeabilized monolayers, the extracellular spores were marked with 1:50 anti–*C. difficile* spore goat serum (recognize specifically to *C. difficile* spores in infections of IECs infection assays) in PBS–1% BSA 1 h RT and 1:400 anti-goat conjugated with CFL 488 secondary antibody (green) (SC362255, Santa Cruz Biotechnologies, USA) in PBS–1% BSA for 1 h RT. The samples were washed three times with PBS and once with sterile distilled water. Samples were then dried at RT for 30 min, and coverslips were mounted using Dako Fluorescence Mounting Medium (Dako, Denmark) and sealed with nail polish. Samples were observed on an Olympus BX53 fluorescence microscope with UPLFLN 100× oil objective (numerical aperture 1.30). Images were captured with the microscope camera for fluorescence imaging Qimaging R6 Retiga and pictures were analyzed with ImageJ (NIH, USA). Extracellular spores or adhered were considerate as spores in phase contrast that were marked in fluorescence. Internalized spores were considered as spores visible in phase contrast, but that does not have fluorescence. A total of ∼300 spores were analyzed per experimental condition.

### Mice used

6-8 weeks C57BL/6 (male or female) were obtained from the breeding colony at the Biological Science Department of Andrés Bello University derived from Jackson Laboratories. Mice were housed with ad libitum access to food and water. Bedding and cages were autoclaved, and mice had a 12-hour cycle of light and darkness. All procedures were performed following the approved protocols by the Institutional Animal Care and Use Committee of the Universidad Andrés Bello.

### Colonic and ileal loop assay

C57BL/6 mice were anesthetized in an isoflurane chamber (RWD USA) with 4% vol vol^−1^ isoflurane (Baxter USA) at and were maintained with 2% vol vol^−1^ during the surgery administrated by air. The intestinal loop model was performed as previously described^18^. Briefly, a midline laparotomy was performed, making 1–cm incision in the abdomen, 1.5 cm ileal, and proximal colon (at 1.0 – 1.5 cm from the cecum as a reference) were ligated with silk surgical suture. To evaluate the effect of NYS or RGD peptide in *C. difficile* spore internalization, mice were treated with 17,000 UI kg^−1^ NYS (*n* = 4) 24h before the surgery. In the loop, as control, mice were treated with 0.9% NaCl (saline) (*n* = 4) then, ligated loops were injected with 3 × 10^8^ *C. difficile* R20291. In the case of RGD, ligated loops were injected with 250 nmol of RGD peptide (*n* = 4). To evaluate the role of BclA3 protein in *C. difficile* spore internalization, the ligated loops were injected with 100 μL of 0.9% weight vol^−1^ NaCl containing 5 × 10^8^ wild-type spore (Δ*pyrE*/*pyrE*+) (SI *n* = 12; colon *n* = 10), Δ*bclA3* (SI *n* = 12; colon *n* = 12) and Δ*bclA3/bclA3+* (SI *n* = 12; colon *n* = 11). The intestine was returned to the abdomen, and the incision was closed. Animals were allowed to regain consciousness. Mice were kept for 5 h and were euthanized. The ligated loops were removed and washed gently in PBS prior to immunostaining, as described below.

### Immunostaining of ileal and colonic loops

First, extracted washed intestinal tissues from the ileal and colonic loops were longitudinally cut, then washed by immersion in PBS 3 times at room temperature (RT). For better visualization of the tissues, they were fixed flat at RT. To perform this, tissues were fixed over a filter paper imbibed with 30% sucrose (Winkler, Chile) in PBS–4% paraformaldehyde (Merck, USA) for at least 15 min. Tissues were transferred to a microcentrifuge tube with the same fixing solution and were incubated at 4 °C overnight. Since fixation of mucus with cross-linking agents, such as paraformaldehyde, cause mucus layer of colon to collapse and shrink to a very tiny lining the epithelia^55^, we did not observe mucus layer in our ileal and colonic loops. Prior to immunostaining, the intestinal and colonic tissues were cut into ∼5 × 5 mm fragments.

To quantify *C. difficile* spore adherence and internalization in the colonic and in the ileal mucosa, tissues were made permeable by incubation with PBS–0.2% Triton X-100 (Merck, USA) and blocked with PBS–3% BSA (Sigma–Aldrich, USA) for 3 h at RT, the same buffer was used for subsequent incubation with antibodies. Tissue was incubated with a primary polyclonal 1:1,000 anti-*C. difficile* spore IgY batch 7246 antibodies (Aveslab USA) in PBS–3% BSA that does not immunoreacted with epitopes of vegetative cells neither with murine microbiota^8^ and with 1:50 phalloidin Alexa-Fluor 568 (#ab176753 Abcam, USA) in PBS–3% BSA overnight at 4 °C to stain the actin cytoskeleton. Following PBS washed, samples were incubated with 1:400 goat anti-chicken IgY secondary antibodies Alexa-Fluor 488 (#ab150173 Abcam USA) in PBS–3% BSA at RT, washed 3 times with PBS and the cellular nuclei stained with 1:1,000 of Hoechst (ThermoFisher, USA) for 15 min at RT.

To perform double immunostaining acc Fn, acc Vn, acc Muc2, with acc Ecad in healthy colonic tissue or acc Muc2 with acc Ecad in the intestinal tissue, first the proximal colon and ilium of 2 independent healthy C57BL/6 mice of 8 weeks old were removed, and mice were sacrificed. Next, tissues were washed by immersion 3 times in PBS at RT and they were fixed flat with 30% sucrose in PBS–4% paraformaldehyde as was described above. Subsequently, tissued cut into ∼5 × 5 mm fragments, then were blocked with PBS-3% BSA (Sigma–Aldrich, USA) for 3 h at RT. And to immunostaining, the acc Ecad, 3 colonic and 1 intestinal tissue fragments of each mouse were incubated with a primary polyclonal 1:200 rat anti-E-cadherin (#ab11512; Abcam USA) in PBS–3% BSA for overnight at 4 °C, then tissues were washed, with PBS, and incubated with 1:300 goat anti-rat IgG secondary antibodies Alexa-Fluor 488 (#A-21470, ThermoFisher, USA) in PBS– 3% BSA for 3 h at RT. Subsequently, to perform the second stain Fn, Vn or Muc2 in tissues stained for accessible Ecad, all tissues fragments were washed with PBS and then for i) stain accessible Fn one colonic tissue fragment of each mice was incubated with 1:200 of rabbit anti-fibronectin (SC9068, Santa Cruz Biotechnologies, USA), or to ii) stain accessible Vn one colonic tissue fragment of each mice was incubated with 1:200 of rabbit anti-vitronectin (SC15332, Santa Cruz Biotechnologies, USA), and finally iii) to stain accessible Muc2 in colonic and ileum tissue one tissue fragment of each mice was incubated with 1:200 of rabbit anti-muc2 ab90007 Abcam (#ab90007. Abcam, USA), in PBS–3% BSA for overnight at 4 °C. at the next day, tissues were incubated with 1:300 of donkey anti-rabbit IgG secondary antibodies Alexa-Fluor 568 (A11036, Invitrogen, USA) for 3 h at RT. Subsequently, tissues were washed and were made permeable by incubation with PBS–0.2% Triton X-100 (Merck, USA) for 1 h at RT. Finally, tissues were incubated with 1:100 phalloidin Alexa-Fluor 647 (A22287 Invitrogen, USA) for 90 min at RT.

The aforementioned immune-stained tissues were subsequently mounted with the luminal side-up. For this, the colonic crypts and the intestinal villi were identified under light microscopy with 10× or 40× magnification and were oriented side-up towards the coverslip. The tissue segment was placed over 5µL of Dako fluorescent mounting medium (Dako, Denmark) applied onto a glass slide, and the tissue covered with 15 µL Dako fluorescent mounting medium and closed with a coverslip. Coverslips were affirmed to the glass slide with vinyl tape to hold the tissue sections in place and were allowed to cure for at least 24 h before imaging.

### Confocal microscopic analysis of ileal and colonic loops

To acquiring images, two confocal fluorescent microscopes were used; a Leica TCS LSI and a Leica SP8 (Leica, Germany) at the Confocal Microscopy Core Facility of the Universidad Andrés Bello. In the fist instance to observe spore internalization in the healthy ileum and colonic mucosa, a Leica TCS LSI was used, with 63× ACS APO oil objective numerical aperture 1.3, and 5× (optical zoom 20×), numerical aperture 0.5. Confocal micrographs were acquired using excitation wavelengths of 405 nm, 488 nm, and 532 nm, and signals were detected with an ultra-high dynamic PMT spectral detector (430–750 nm). Emitted fluorescence was split with four dichroic mirrors (QD405 nm, 488 nm 561 nm, and 635 nm). Images (1,024 × 1,024 pixels). To observe the sites with acc Fn and Vn in the intestinal barrier and to evaluate the adherence and internalization of the Δ*bclA3* spore mutant to the intestinal barrier or to evaluate spore adherence and internalization in mice treated with RGD and NYS confocal images were acquired in Leica SP8 was used with HPL APO CS2 40× oil, numerical aperture 1.30. Signals, 3 PMT spectral detector PMT1 (410-483) DAPI PMT2 (505-550) Alexa-Fluor 488 PMT3 (587-726) Alexa-Fluor 555. Emitted fluorescence was split with dichroic mirrors DD488/552. Three-dimensional reconstructions of intestinal epithelium were performed using ImageJ software (NIH, USA). Villi and crypts were visualized by Hoechst and phalloidin signals. 3D reconstruction videos were performed with Leica software LASX 3D (Leica, Germany).

To quantify cells of the colonic and ileum mucosa with accessible proteins immunodetected, confocal images with a 1–µm Z step size were filtered with Gaussian Blur 3D (sigma x: 0.6; y: 0.6; z:0.6) and quantifies with cell counting plug-in of ImageJ 1,000 - 1,200 cells were counted in an area 84,628µm^2^ per mice.

To quantify spore adherence and internalization, confocal images with a 0.7–µm Z step size were analyzed. Adhered spores were considered as fluorescent-spots that were in narrow contact with the actin cytoskeleton (visualized with phalloidin), and internalized spores were considered as fluorescent-spots that were inside the actin cytoskeleton in the three spatial planes (orthogonal view)^23,56^. The analyzed area for each tissue was 338,512 µm^2^ per animal.

Then we evaluated the spore distribution of adhered and internalized in the colonic and in the intestinal mucosa. To measure the distribution of adhered spores in the colonic and ileum mucosa, the perpendicular distance from the center of the spore to the epithelium was measured using ImageJ (NHI, USA). In the case of internalized spores, we measured the perpendicular distance from the center of the spore to the mucosa surface or from the closest crypt membrane. For ileum mucosa, we measure the perpendicular distance from the center of the spore to the villus tip or to the villus membrane.

### Visualization of spore internalization in intestinal epithelial cells *in vitro* by confocal microscopy

Differentiated Caco-2 cells, and T84 cells cultured onto Transwell (Corning USA) until 1,000**–**2,000 Ω. Cells were infected for 5 h with an MOI of 10 with *C. difficile* spores previously stained with Alexa Fluor 488 Protein Labeling Kit (Molecular Probes, USA) according to the manufacturer’s instruction. Cells were washed 2 twice with PBS and were permeabilized with PBS–0.06%-Triton X-100 (Merck, USA) for 10 min at RT, were washed and incubated with 1:150 phalloidin Alexa-Fluor 568 (#ab176753 Abcam, USA) in PBS–1% BSA for 1h at RT. Then cells were washed, fixed and visualized in a confocal microscopy Olympus FV1000 of the Confocal Microscopy Core Facility of the Andrés Bello University.

### Sample preparation for Transmission Electron Microscopy and Immuno-electron microscopy

To visualize internalized spores in IECs, six-well plates containing differentiated Caco-2 cells or T84 cells cultured in transwell as was described above were infected for 5 h at 37 °C at an MOI of 20 with *C. difficile* R20291 spores pre-incubated 1h at 37 °C with 100 µL of NHS (Complement Technology USA) for each well, and then was suspended in the infection volume of 1mL; FBS final concentration 10% vol vol^−1^ FBS. Unbound spores rinsed o□, and cells were scraped, fixed, and processed, as is described below.

To evaluate the binding of Fn and Vn to the surface of *C. difficile* spores, 4 × 10^7^ *C. difficile* spore-were incubated in PBS–0.2% BSA containing 10 µg ml^−1^ of human Fn and Vn for 1h at 37 °C. The spores were then washed three times (18,400×*g* for 5 minutes at RT) with PBS. Then the spores were pelleted by one cycle of centrifugation at 18,400 × *g* for 10 min. Pellets were then resuspended in 200 µl PBS-1% BSA, incubated for 30 min at RT, and then sedimented by centrifugation at 18,400 × *g* for 10 min at RT. Pellets were resuspended as above and incubated with primary antibody 1:200 rabbit pAb against Fn (SC9068, Santa Cruz Biotechnology USA) or Vn (SC15332, Santa Cruz Biotechnology, USA) in PBS-BSA 1% for 1 h at RT. The excess of antibody was eliminated by three cycles of centrifugation at 18,400×*g* for 5 min at RT and resuspension in PBS–0.1% BSA. Spore suspensions were then incubated for 1 h with 1:20 donkey anti-rabbit IgG antibody coupled to 12–nm gold particles (Abcam ab105295, USA) in PBS-1% BSA for 1h at RT. And were washed by triple centrifugation at 18,400×*g* for 5 min. Subsequently, samples were fixed and processed, as is described below.

To visualizes if the collagen-like BclA3 exosporium protein forms the hair like-extension of *C. difficile* spores, ∼2×10^8^ *C. difficile* purified spores of wild-type R20291 (Δ*pyrE*/*pyrE*^+^), Δ*bclA3* and Δ*bclA3*/*bclA3*^+^ fixed and processes as is described here below.

### Sample processing and staining for transmission electron microscopy

The aforementioned spore or monolayers of infected IECs samples were fixed with freshly prepared with 2.5% glutaraldehyde 1% paraformaldehyde in 0.1 M cacodylate buffer (pH 7.2) overnight at 4 °C, rinsed in cacodylate buffer, and stained for 30 min with 1% tannic acid. Then samples were serially dehydrated with acetone 30% (with or without 2% uranyl acetate) for 20 min, 50% for 20 min, 75% for 20 min, 90% for 20 min, and twice with 100% for 20 min, embedded in spurs resin at ratio acetone: spurs of 3:1, 1:1, and 1:3 for 40 min each and then resuspended in spurs for 4 h and baked overnight at 65 °C, and prepared for transmission electron microscopy as previously described^15^. Thin sections (90 nm) obtained with a microtome were placed on glow discharge carbon-coated grids for negative staining and double lead stained with 2% uranyl acetate and lead citrate. Grids were analyzed with a Phillips Tecnai 12 Bio Twin electron microscope of the Universidad Católica de Chile.

### R-CDI mouse model

Antibiotic cocktail (ATB cocktail) was administrated, as was previously described ^8^. Briefly, an antibiotic cocktail containing 40 mg kg^−1^ kanamycin (Sigma–Aldrich, USA), 3.5 mg kg^−^1 gentamicin (Sigma–Aldrich, USA), 4.2 mg kg^−1^ colistin (Sigma–Aldrich, USA), 21.5 mg kg^−^1 metronidazole (Sigma–Aldrich, USA) and 4.5 mg kg^−^1 vancomycin (VAN) (Sigma–Aldrich, USA) was administrated via gavage for 3 days (days -6 to -4 before the infection). Then 1 day before the infection (day -1), an intraperitoneal (i.p.) injection of 10 mg kg^−1^ clindamycin (Sigma–Aldrich, USA) was administrated to all mice. The next day all mice were infected via gavage with 1 × 10^7^ spores R20291, and on day 3 post-infection, DPBS containing 17,000 UI kg^−1^ of nystatin and 50 mg kg^−1^ vancomycin or vancomycin alone (as control) was orally administered for 5 days to 18 and 23 mice, respectively.

To evaluate the role of the exosporium protein BclA3 in the R-CDI, 40 treated mice with antibiotic cocktail followed with clindamycin, were infected orally with 100 µl PBS containing 5 × 10^7^ *C. difficil*e spore strain R20291 of wild-type (Δ*pyrE*/*pyrE*+) (*n* = 10), Δ*bclA3* (*n* = 16) or Δ*bclA3/bclA3*+ (*n* = 14) strains. *C. difficile*-infected mice were housed individually in sterile cages with *ad libitum* access to food and water. All procedures and mouse handling were performed aseptically in a biosafety cabinet to contain spore mediated transmission and cross-contamination. Mice were daily monitored for loss weight, aspect, and diarrhea were measured to determine the endpoint of each animal.

During the entire experiment, the clinical condition (sickness behaviors and fecal samples) of mice was monitored daily with a scoring system. The presence of diarrhea was classified according to severity as follows: (i) normal stool (score = 1); (ii) color change and/or consistency (score = 2); (iii) presence of wet tail or mucosa (score = 3); (iv) liquid stools (score = 4). A score higher than 1 was considered as diarrhea^57^. At the end of the assay, animals were sacrificed with a lethal dose of ketamine and xylazine. Cecum content and colonic tissues were collected.

### Quantification of *C. difficile* spores from feces and colon of mice

To quantify *C. difficile* spores in feces, daily collected fecal samples were stored at −20 °C until spore quantification.

Feces were hydrated in 500 µL sterile Milli-Q water overnight at 4 °C and then mixed with 500 µL of absolute ethanol (Merck, USA) for 60 min at RT. Then, serially dilutions of the sample were plated onto selective medium supplemented with 0.1% weight vol^−1^ taurocholate, 16 µg ml^−1^ cefoxitin, 250 µg mL^−1^ L-cycloserine and 1.5% weight vol^−1^ (BD, USA) (TCCFA plates). The plates were incubated anaerobically at 37 °C for 48 h, colonies counted, and results expressed as the Log_10_ (CFU g^−1^ of feces)^58^. Colonic tissue collected at the end of the experiment was washed three times with PBS. The tissue *C. difficile* spore-load was determined in the proximal colon, medium colon, distal colon, and cecum tissue. Tissues were weighed and adjusted at a concentration of 100 mg mL^−1^ with a 1:1 mix of PBS: absolute ethanol, then homogenized and incubated for 1 h at RT. The amounts of viable spores were quantified plating the homogenized tissue onto TCCFA plates, as described previously^18^. The plates were incubated anaerobically at for 48 h at 37 °C. Finally, the colony count was expressed as the Log_10_ (CFU g^−1^ of the tissue).

### Cecum content cytotoxicity assay in Vero cells of infected mice during R-CDI

Vero cell cytotoxicity was performed as described previously^59^. At first, 96-well flat-bottom microtiter plates were seeded with Vero cells at a density of 10^5^ cells well^−1^. Mice cecum contents were suspended in PBS at a ratio of 1:10 (100 mg mL^−1^ of cecum content), vortexed and centrifuged at 18,400×*g* for 5 min, the supernatant was sterilized with a 0.22µm filter and serially diluted in DMEM supplemented with 10% vol vol^−1^ FBS and 100 U ml^−1^ penicillin, and 100 µg ml^−1^ streptomycin; then 100 µL of each dilution was added to wells containing Vero cells. Plates were screened for cell rounding 16 h after incubation at 37 °C. The cytotoxic titer was defined as the reciprocal of the highest dilution that produced rounding in at least 80% of Vero cells per gram of luminal samples under 20× magnification.

## Statistical analysis

Prism 7 (GraphPad Software, Inc.) was used for statistical analysis. Student’s *t*-test and the nonparametric test was used for pairwise comparison. Significance between groups was done by Mann-Whitney unpaired *t*-test. Comparative study between groups for *in vitro* experiments was analyzed by analysis of variance with post-hoc Student *t*-tests with Bonferroni corrections for multiple comparisons, as appropriate. A *P*-value of ≤ 0.05 was accepted as the level of statistical significance. Differences in the percentages of mice with normal stools, as well as percentages of mice with *C. difficile* infection, were determined by Gehan-Breslow-Wilcoxon test.

## Acknowledgments

This work was funded by FONDECYT Regular 1191601 and FONDECYT Regular 1151025 to D.P-S. Millennium Nucleus of the Biology of the Intestinal Microbiota to D.P-S. PC-C had been supported by ANID doctoral fellowship 21161395 (Chile) and JO-A by OAICE-91-2018 of Universidad de Costa Rica. The authors acknowledge Rosario Hernandez-Armengol for technical assistance and Miriam Barros for useful discussion on image processing at the Confocal Microscopy Core Facility of the Universidad Andrés Bello. We certify that funding sources had no implication in the study design, collection of data, analysis, and interpretation of data.

## Extended Data Figure Legends

**Extended Data Fig. 1** | **Adherence of *C. difficile* spores to the colonic mucosa**. *z*-plane and orthogonal view of confocal micrographs of fixed whole-mount colonic tissue of mice. Panels **a**−**c, g** micrographs of colon of C57BL/6 mice infected in a colonic loop model with 5 × 10^8^ *C. difficile* R20291 spores for 5 h. *C. difficile* spores are shown in red, F-actin is shown in green and nuclei in blue (fluorophores colors were digitally reassigned for a better representation). Panels **d**−**f, h** are a magnification of panels **a**−**c, g** respectively. Highlighted are *C. difficile* spores close the apical membrane of cells. Micrographs are representative of 3 independent mice. Bars **a**−**c, g** 50 μm, **d**−**f, h** 10 μm.

**Extended Data Fig. 2** | **Adherence of *C. difficile* spores to the intestinal mucosa**. *z*-plane and orthogonal view of confocal micrographs of fixed whole-mount colonic tissue of mice. Panels **a**−**c, g** micrographs of ileum mucosa of C57BL/6 mice infected in an ileal loop model with 5 × 10^8^ spores *C. difficile* R20291 spores for 5 h. *C. difficile* spores are shown in red, F-actin is shown in green and nuclei in blue (fluorophores colors were digitally reassigned for a better representation). Panels **d**−**f, h** are a magnification of panels **a**−**c, g** respectively. Highlighted are *C. difficile* spores close the apical membrane of cells. Micrographs are representative of 3 independent mice. Bars **a**−**c, g**, 50 μm, **d**−**f, h**, 10 μm.

**Extended Data Fig. 3** | **Internalization of *C. difficile* spores to the colonic mucosa *in vivo***. *z*-plane and orthogonal view of confocal micrographs of fixed whole-mount colonic tissue of mice. Panels **a**−**c, g**, Confocal micrographs of the colonic mucosa of C57BL/6 mice infected in a colonic loop model with 5 × 10^8^ *C. difficile* R20291 spores for 5 h. *C. difficile* spores are shown in red, F-actin is shown in green and nuclei in blue (fluorophores colors were digitally reassigned for a better representation). Panels **d**−**f, h** are a magnification of panels **a**−**c, g**, respectively. Highlighted are *C. difficile* spores close the apical membrane of cells. Micrographs are representative of 3 independent mice. Bars **a**−**c, g**, 50 μm, **d**−**f, h**, 10 μm.

**Extended Data Fig. 4** | **Internalization of *C. difficile* spores into the intestinal mucosa**. *z*-plane and orthogonal view of confocal micrographs of fixed whole-mount colonic tissue of mice. Panels **a**−**c** micrographs of the ileum mucosa of C57BL/6 mice infected in an ileal loop model with 5 × 10^8^ spores *C. difficile* R20291 spores for 5 h. *C. difficile* spores are shown in red, F-actin is shown in green and nuclei in blue (fluorophores colors were digitally reassigned for a better representation). Panels **d**−**f, h** are magnification of panels **a**−**c, g** respectively. Highlighted are *C. difficile* spores close the apical membrane of cells. Micrographs are representative of 3 independent mice. Bars **a**−**c, g** 50 μm, **d**−**f, h** 10 μm.

**Extended Data Fig. 5** | **Distribution of internalized *C. difficile* spore in colonic and in ileum mucosa. a** Schematics of the method applied for measurements of distances of internalized spores in the ileum mucosa. Distribution of the distance of internalized *C. difficile* spores to the **b** villus tip or to **c** the villus membrane. **d** Schematics of the method applied for measurements of distances of internalized spores in the colonic mucosa. **e** Distribution of the distance of internalized *C. difficile* spores to the colonic epithelium surface or **f** to the closest crypt axis. In the ileum, distance from 34 internalized *C. difficile* spores to the villus tip and the distance of 54 internalized *C. difficile* spores to the villus membrane is shown. In the colon, the distance of 61 internalized *C. difficile* spores was evaluated.

**Extended Data Fig. 6** | **Confocal microscopy of internalized *C. difficile* spores into intestinal epithelial cells *in vitro***. *z*-plane and orthogonal view of confocal micrographs of **a, b** polarized T84 cells infected with NHS-treated R20291 spores for 5 h. **c, d** Intestinal epithelial Caco-2 cells infected with NHS-treated *C. difficile* R20291 spores. *C. difficile* spores are shown in red, F-actin is shown in green, (fluorophores colors were digitally reassigned for a better representation). Scale bar, 10 µm.

**Extended Data Fig. 7** | **Internalized spores are not stained with anti-*C. difficile* spore goat serum in non-permeabilized cells, and serum increases the internalization into Caco-2 cells. a**, Cells were infected for 3 h with FBS treated *C. difficile* R20291 spores, and without permeabilization, spores were immunodetected as is described in methods. Bright spores that were not detected with fluorescent-labeled antibodies were considered as internalized. Dynamic of *C. difficile* spore-entry of spores pre-incubated with FBS in infected **i** Caco-2, and **j** T84 cells for 3h in the presence of 10% FBS. **k** Internalization of *C. difficile* spores of clinical isolates of various ribotypes into Caco-2 cells. Error bars indicate the mean ± S.E.M. Statistical analysis was performed by Student’s *t*-test. ns, indicates non-significant differences, \**P* < 0.01.

**Extended Data Fig. 8** | **The spore entry into Caco-2 cell line in the presence of normal human serum is inhibited by RGD peptide. a**−**b** Caco-2 cells monolayer differentiated, and **c–f** undifferentiated. **a, c, e** show relative internalization while **b, d, f** shows relative adherence compared to the control (0 μg mL^−1^ RGD). Before the infection, spores were incubated with **a, b, e, f** culture media; **c, d** with NHS. Cells were incubated with 0 and 5 µg mL^−1^ of RGD peptide and infected with R20291 *C. difficile* spores pre-incubated with DMEM (without serum) or NHS. The data represents the mean of three biological experiments. Error bars indicate the mean ± S.E.M. Statistical analysis was performed by Student’s *t*-test, ns, indicates non-significant differences, \**P* < 0.0001.

**Extended Data Fig. 9** | ***C. difficile* spore entry of *C. difficile* spore pre-incubated Fn and Vn and Caco-2 cells pre-incubated Fn and Vn. a**−**d** Caco-2 cells monolayers undifferentiated and **e, f** 8 days differentiated. **a, c, e** shows relative internalization, while **b d, f** shows relative adherence compared to the control (DMEM). Before infecting cells, **a, b** spores or **c, d** Caco-2 cells were pre-treated with NHS, Fn or Vn, and subsequently infected with *C. difficile* R20291 spores. The data represents the mean of three biological experiments. Error bars indicate the mean ± S.E.M. Statistical analysis was performed by Student’s *t*-test, ns indicates non-significant differences \**P* < 0.005; \*\**P* < 0.001.

**Extended Data Fig. 10** | **Accessibility Muc2 in the small intestine. a** Confocal micrographs of fixed whole-mounted healthy colon of C57BL/6 mice for acc Muc2 with acc Ecad. The main figure shown a 3D projection, below magnifications and a z-stack of representative cells with the different immunostaining. **b** Shown the cell repartition of cells immunodetected for acc Ecad. **c** Cell repartition of cells immunodetected for Muc2. **d** Cell repartition of total acc Muc2 cells that were immunodetected for acc E-cad. Acc-Muc2 is shown in green Acc Ecad is shown in red and F-actin in grey (fluorophores colors were digitally reassigned for a better representation). Scale bar, 20µm, in magnifications 10µm. Micrographs are representative of mice (*n* = 2). 1,000 - 1,200 cells were counted for each mouse in an area 84,628µm^2^. Error bars indicate mean ± S.E.M.

**Extended Data Fig. 11** | **The spore entry into Caco-2 cells mediated by Fn occurs through integrin subunits** α**5 and** β**1; while internalization mediated Vn is through integrin subunits** α**v and** β**1. a**−**f** undifferentiated and **g, h** differentiated Caco-2 cell monolayer, and were pre-incubated with **a**−**d** 1, 3 or 5 µg mL^−1^ of RGD peptide or **e**−**h** 10 µg mL^−1^ of antibody against α_v_, α_2,_ α_5_, β_1_, β_3_, non-immune IgG antibody in DMEM and infected with *C. difficile* R20291 spores pre-incubated with 10 µg mL^−1^ **a, b, e, f** Fn, or **c, d, g, h** Vn. **a, c, e, g** Show relative internalization while **b, d, f, h** relative adherence compared to the control (no RGD or non-immune IgG antibody). Error bars indicate the mean ± S.E.M. Statistical analysis was performed by Student’s *t*-test, ns indicates non-significant differences, *P* < 0.01, \*\**P* < 0.001.

**Extended Data Fig. 12** | **Serum-free medium does not promote internalization of *C. difficile* spores dependent of integrins** α_**v**,_ α_**5**_, **and** β_**1**_. **a** Internalization or **b** adherence of *C. difficile* pre-treated 1 h with DMEM in CHO cells ectopically expressing α_v_, α_5_, β_1_ integrins relative to the control (wild-type CHO cells). Error bars indicate the mean ± S.E.M. Statistical analysis was performed by Student’s *t*-test, ns indicates non-significant differences, *P* < 0.01, \*\**P* < 0.001.

**Extended Data Fig. 13** | **Quantification of *C. difficile* spores associated with Fn and Vn by TEM. a, b**, percentage of the total of labeled spores treated with Fn or Vn (Fn + or Vn + respectively) and the negative control without protein (Fn - or Vn - as appropriate). Error bars indicate the mean ± S.E.M. Statistical analysis was performed by Student’s *t*-test, ns indicates non-significant differences, \**P* < 0.001.

**Extended Data Fig. 14** | **Deletion of *bclA3* genes in R20291**. The deletion of *bclA3* was done by allelic exchange through a schematic representation of the deletion of *bclA3*, leaving a small peptide in frame. Construction and characterization of *bclA3* mutant in *C. difficile*. **a** Schematic representation of in-frame deletion of *bclA3*. **b** The size of *bclA3* loci was verified by PCR using detection primers (Table S2).

**Extended Data Fig. 15** | **Immunofluorescence intensity of the anti-spore in *bclA3* mutant *C. difficile* spores. a** Wild-type (Δ*pyrE*/*pyrE*^+^), Δ*bclA3* and Δ*bclA3*/*bclA3*^+^ R20291 spores are recognized by anti-*C. difficile* spore goat serum. Δ*pyrE/pyrE*^+^, (wild-type; wt), Δ*bclA3*, and complemented mutant (Δ*bclA3/bclA3*^+^) R20291 spores were fixed on glass coverslips treated with poly-lysine; then the samples were blocked with PBS–1% BSA and were labeled for 1 h with goat-anti-spore serum and incubated with secondary antibody anti-goat CFL 488-conjugated; then, micrograph were captured with epifluorescence microscopy. **b** Quantitative analysis of the fluorescence (Fl.) intensity in spores of Δ*pyrE/pyrE*^+^ (dark line), Δ*bclA3* (red line) and Δ*bclA3/bclA3*^+^ (blue line), the values shown in the graphs the normalized fluorescence intensity from 150 spores of Δ*pyrE/pyrE*^+^, Δ*bclA3* and Δ*bclA3/bclA3*^+^ strain. **c** Distribution of the fluorescence intensity of wild-type (dark bars), Δ*bclA3* (red bars) and Δ*bclA3/bclA3*^+^ (blue bars) spores.

**Extended Data Fig. 16** | **Effect of absence of BclA3 on spore-entry and adherence to non-phagocytic cells**. Relative *C. difficile* spore **a, c** internalization and **b, d** adherence of HeLa cells infected with *C. difficile* spores Δ*pyrE/pyrE*^+^ (wild-type), Δ*bclA3*, and Δ*bclA3/bclA3*^+^ pre-incubated with 10 µg of **a, b** Fn or **c, d** Vn. **e**−**h** shown the relative *C. difficile* spore adherence of CHO cells ectopically expressing integrins; **e**, CHO-α_5_, **f, h** CHO-β_1_ and **g** CHO-α_v_ and infected with *C. difficile* spores Δ*pyrE/ pyrE*^+^ (wild-type), Δ*bclA3*, and Δ*bclA3/bclA3*^+^ pre-incubated with 10 µg of **e, f** Fn or **g, h** Vn. The data represents the mean of three biological experiments. Error bars indicate the mean ± S.E.M. Statistical analysis was performed by Student’s *t*-test, ns indicates non-significant differences, \**P* < 0.001.

**Extended Data Fig. 17** | **Representative confocal micrograph of adhered and internalized *C. difficile* spores in colonic mucosa with** Δ***bclA3* spores and Extended data of BclA exosporium protein contributes to the recurrence of *C. difficile* infection**. There are supplementary figures of the experiment detailed in Fig 6. Confocal micrograph of the whole-mount fixed colon. C57BL/6 mice infected in a colonic loop model with 5 × 108 *C. difficile* R20291 spores. Panels shown confocal micrographs of adhered and internalized spores in colon loop infected with strains **a** wild-type (Δ*pyrE*/*pyrE*^+^), **b** Δ*bclA3*, and **c** Δ*bclA3/bclA3*^+^. The main figure shown a 3D projection, below the z-stack, magnification, and the orthogonal view. *C. difficile* spores are shown in red, F-actin is shown in green and nuclei in blue (fluorophores colors were digitally reassigned for a better representation). The white arrow indicates internalized *C. difficile* spores, empty arrow indicates adhered *C. difficile* spore. Micrograph are representative of mice wild type (*n* = 10); Δ*bclA3* (*n* = 12) and Δ*pyrE*/*pyrE*^+^ (*n* = 11). These are supplementary figures from the experiment of R-CDI in animals infected with Δ*bclA3* spores (Fig. 6). Animals were infected with wild-type (Δ*pyrE*/*pyrE*^+^; *n* = 10), Δ*bclA3* (*n* = 16) and Δ*bclA3*/*bclA3*^+^, (*n* = 14) and were monitored daily for **d** relative weight; **e** time to diarrhea during CDI. **f** *C. difficile* spore CFU in feces during CDI and R-CDI. Spore adherence to the colonic tract was evaluated on day 11 to **g** Cecum, **h** proximal colon, **i** distal colon and, **j** cytotoxicity of the cecal content. Error bars indicate the mean ± S.E.M. Statistical analysis was performed by **d, f**, Kruskal-Wallis, post-Dunn’s test, **e**, Log-rank (Mantel-Cox), **g**−**j**, Mann-Whitney test, ns indicates non-significant differences, * *P* < 0.05, \*\**P* < 0.01, ***P < 0.001. Scare bar, 20µm.

**Extended Data Fig. 18** | **Extended data of Fig 7**. These are supplementary figures from the experiment detailed in Fig 7. *C. difficile* spore **a** internalization and **b** adherence in monolayers of T84; cells were pre-treated with 30 µM of nystatin (shown as NYS) for 1 h and were subsequently infected with *C. difficile* spores R20291 pre-treated for 1 h with FBS. **c** Cellular viability after nystatin treatment was determined by using MTT assay. **d–f** Confocal micrograph of the whole-mount fixed colon. C57BL/6 mice infected in a colonic loop model with 5 × 10^8^ *C. difficile* R20291spores. Panels shown adhered and internalized spores of **d** untreated mice (Ctrl), **e** mice treated 24h before the surgery with nystatin, and administered in the loop in the presence of: **f** 250 nmol of RGD peptide in the loop. The main figure shown a 3D projection, below the z-stack, magnification, and the orthogonal view. *C. difficile* spores are shown in red, F-actin is shown in green and nuclei in blue (fluorophores colors were digitally reassigned for a better representation). White arrow indicates internalized *C. difficile* spores, empty arrow indicates adhered *C. difficile* spore. Micrographs are representative of mice (*n* = 4). These are supplementary figures from the experiment of R-CDI with animals treated with nystatin and RGD. Animals were infected with *C. difficile* R20291 and were monitored daily for **g** weight loss, **h** diarrhea, and **i** *C. difficile* spore CFU in feces during CDI and R-CDI. For mice treated with nystatin and vancomycin (*n* = 18) or vancomycin alone as control (*n* = 23) from days 3-7. Vancomycin is shown as VAN. Error bars indicate the mean ± S.E.M. Statistical analysis was performed by **a, b**, Student’s *t*-test, **f** and **h**, Kruskal-Wallis, post-Dunn’s test, **g**, Log-rank (Mantel-Cox), ns indicates non-significant differences, \**P* < 0.05, \*\**P* < 0.01, \*\*\**P* < 0.001. Scale bar, 20µm.

## Supplementary Videos

**Video 1. Wild-type *C. difficile* spores internalize in colonic mucosa**.

Travel through a confocal *Z*-stack from the apical face of the colonic mucosa. *C. difficile* spore is shown in red, F-actin is shown in green and nuclei in blue (fluorophores colors were digitally reassigned for a better representation). Arrows indicate internalized spores.

**Video 2. Wild-type *C. difficile* spores internalize in the ileum mucosa**.

Travel through a confocal *Z*-stack from the apical face of the ileum mucosa. *C. difficile* spore is shown in red, F-actin is shown in green and nuclei in blue (fluorophores colors were digitally reassigned for a better representation). Arrows indicate internalized spores.

