## Extended Figures for "*Clostridioides difficile* spore-entry into intestinal epithelial cells contributes to recurrence of the disease"

### Extended Data Fig. 1

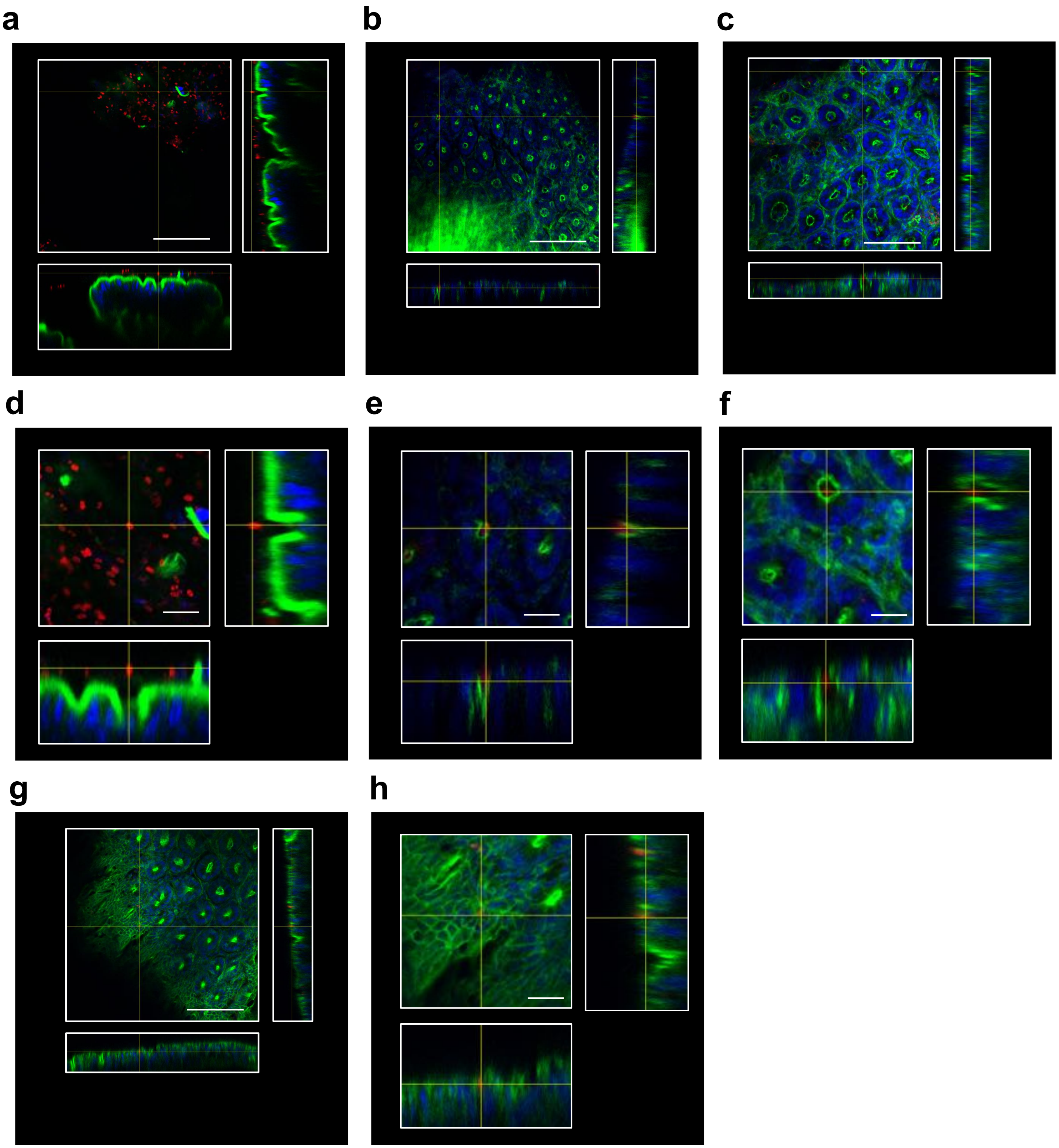

### Extended Data Fig. 2

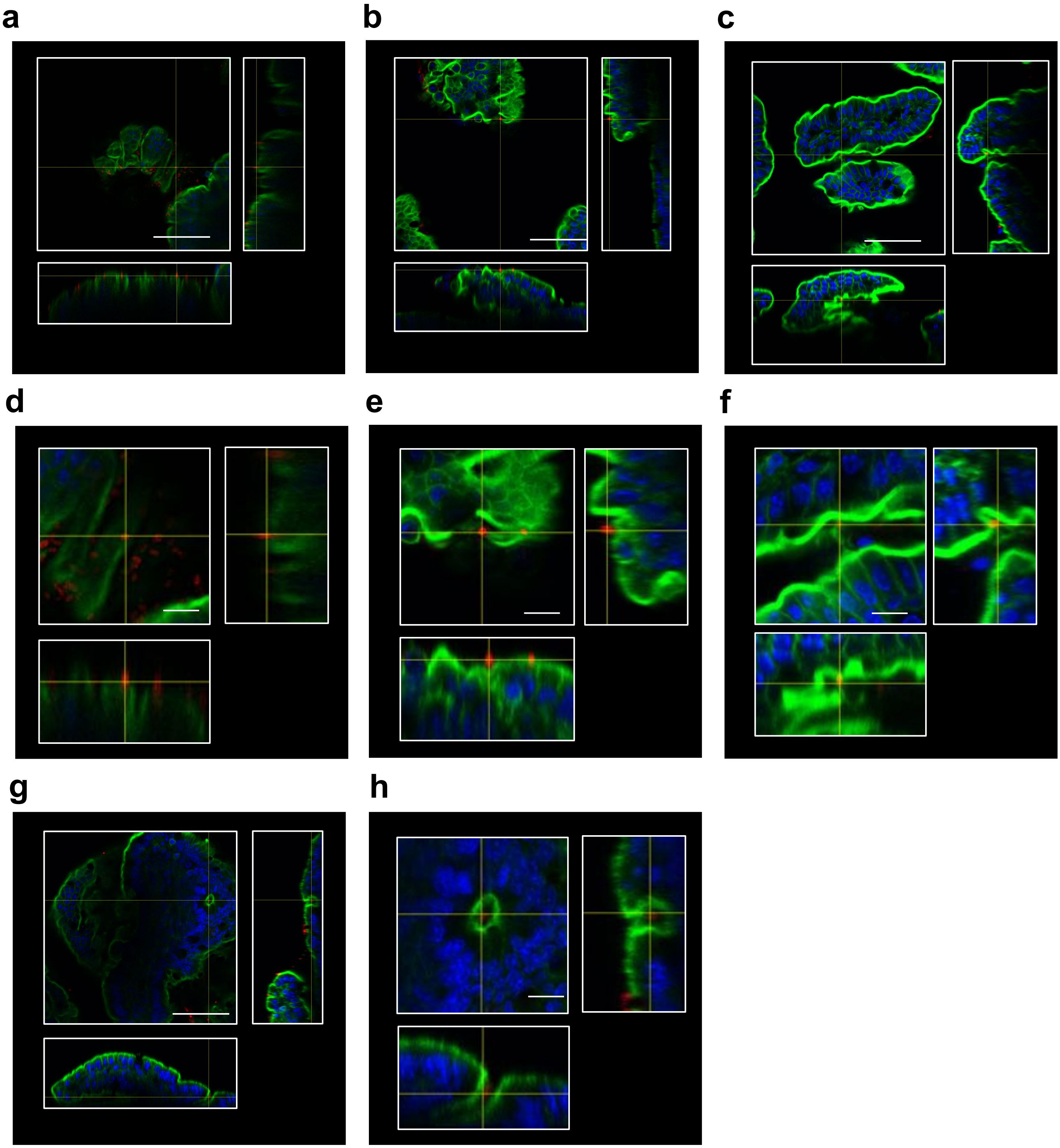

### Extended Data Fig. 3

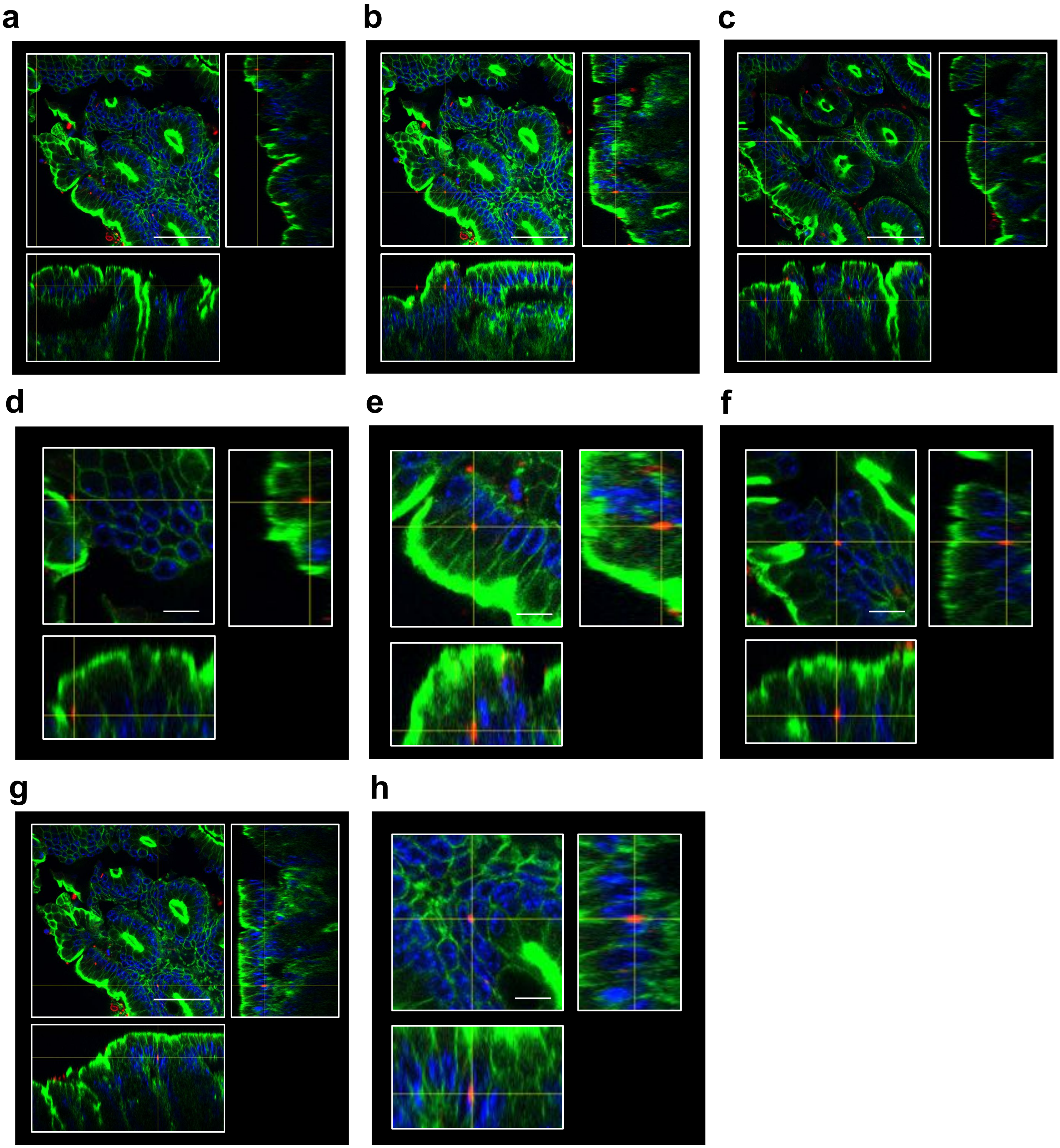

### Video 1 Colon

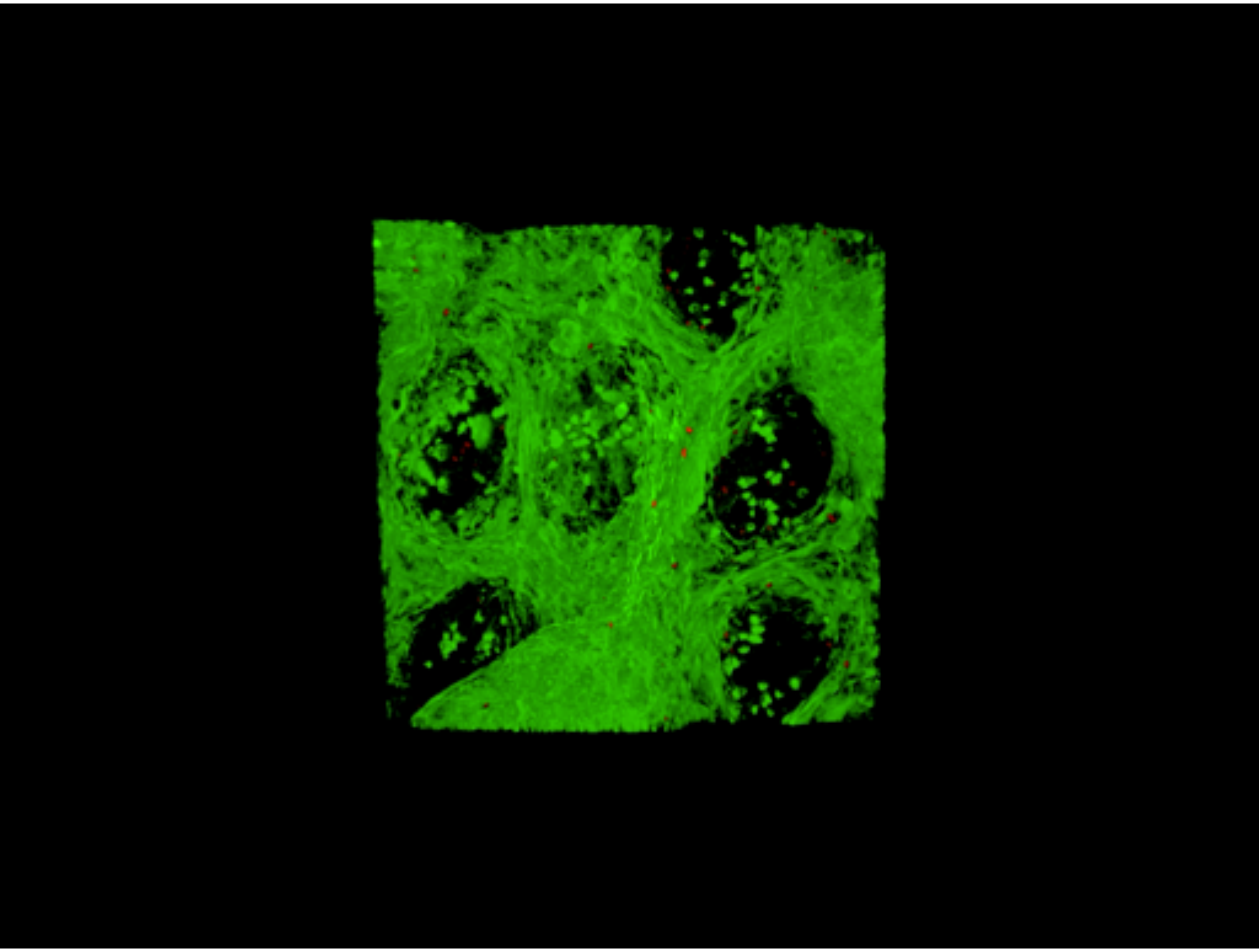

### Extended Data Fig. 4

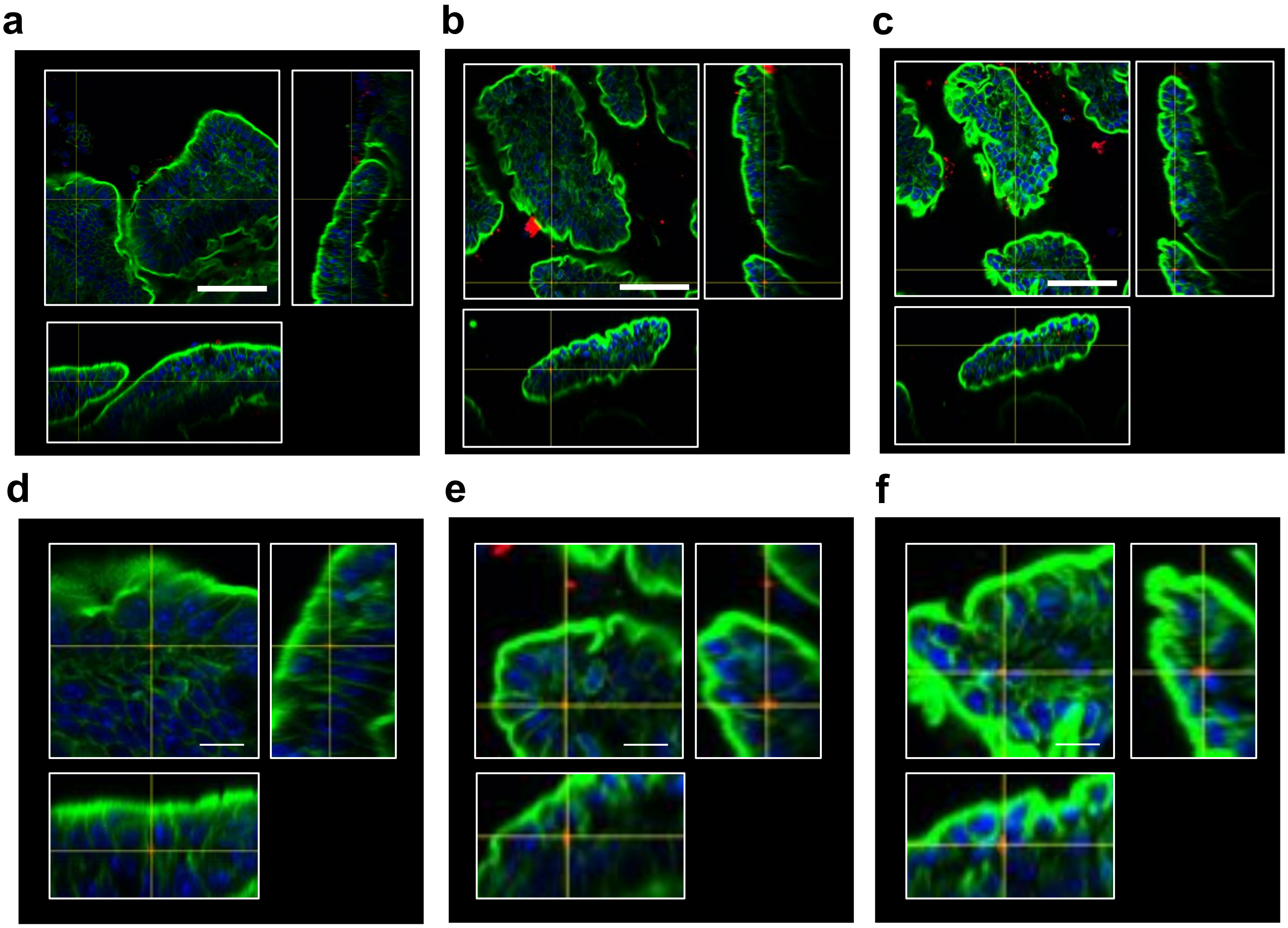

### Video 2 Small intestine

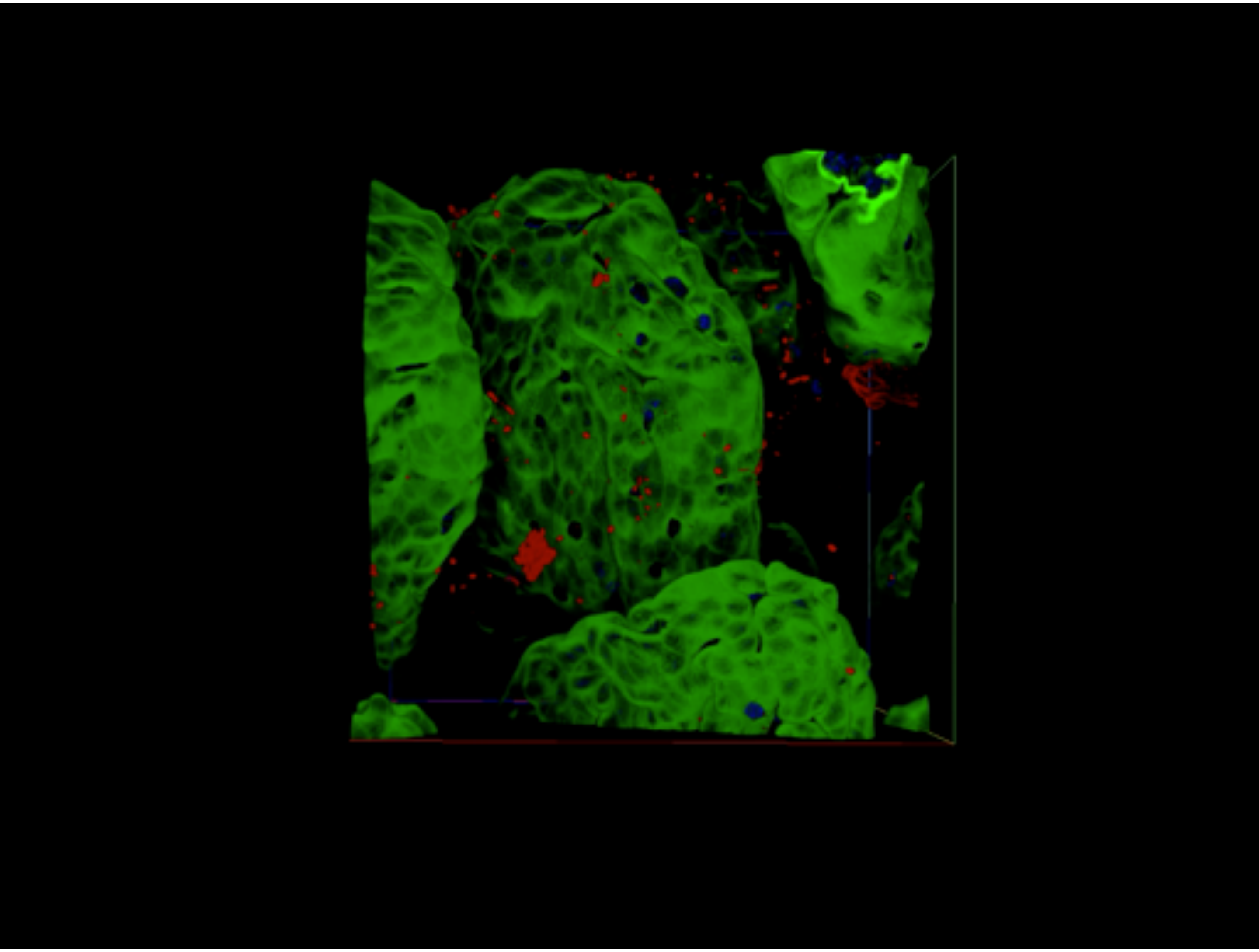

### Extended Data Fig. 5

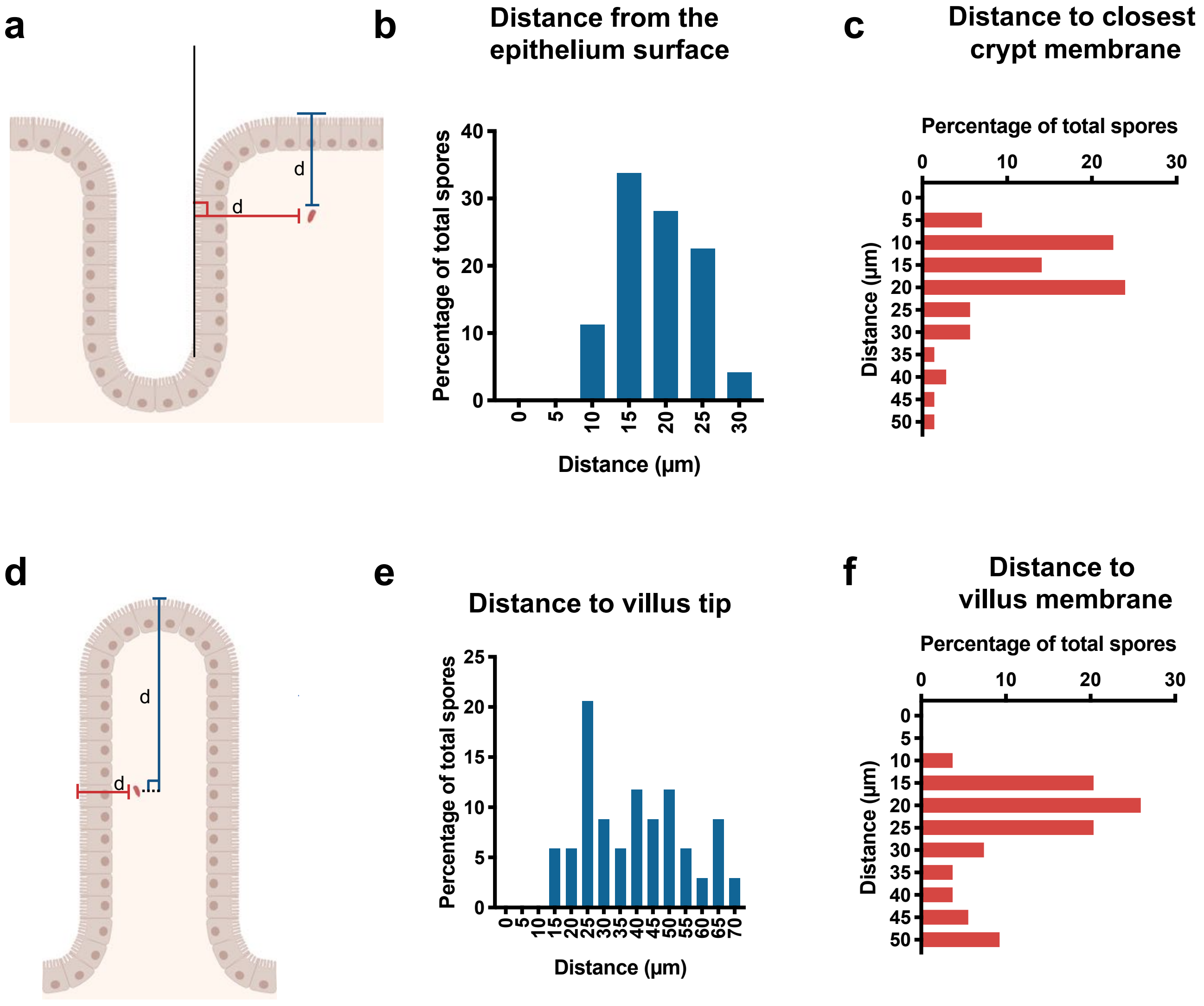

### Extended Data Fig. 6

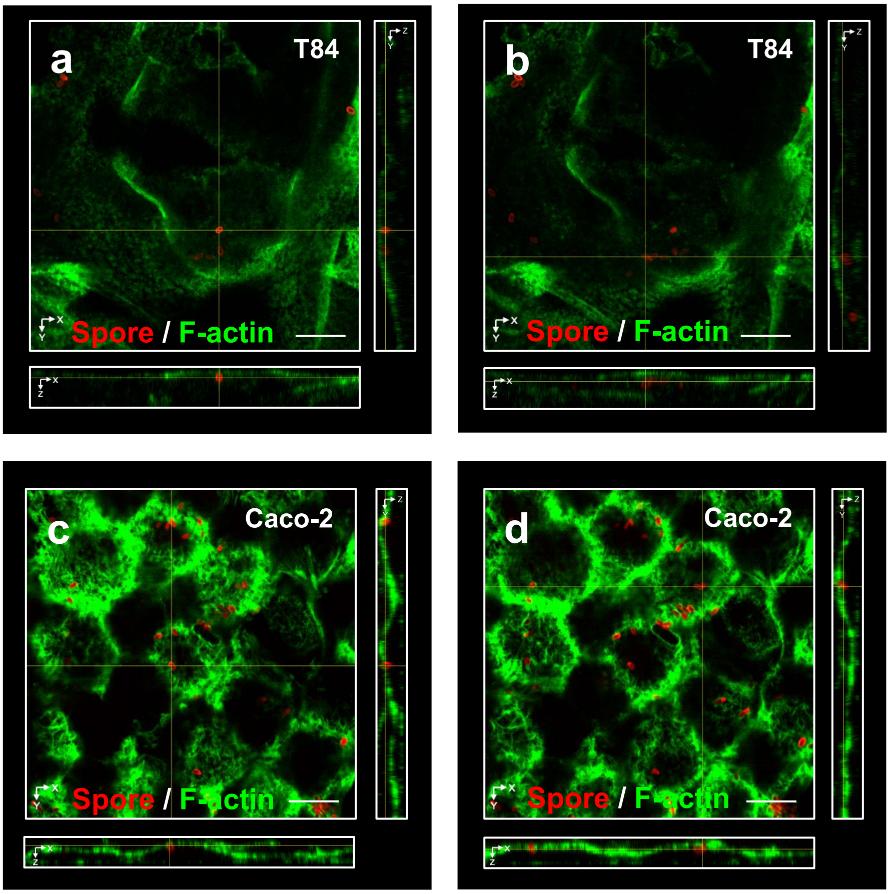

### Extended Data Fig. 7

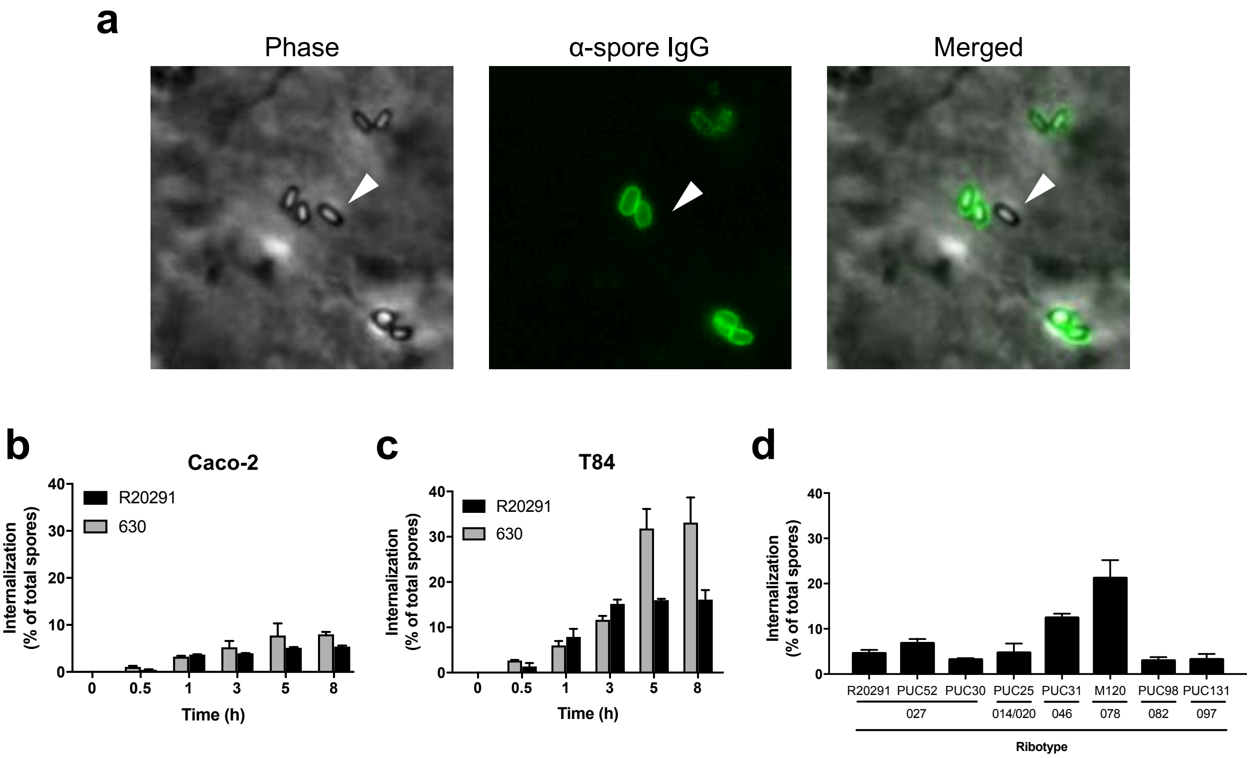

### Extended Data Fig. 8

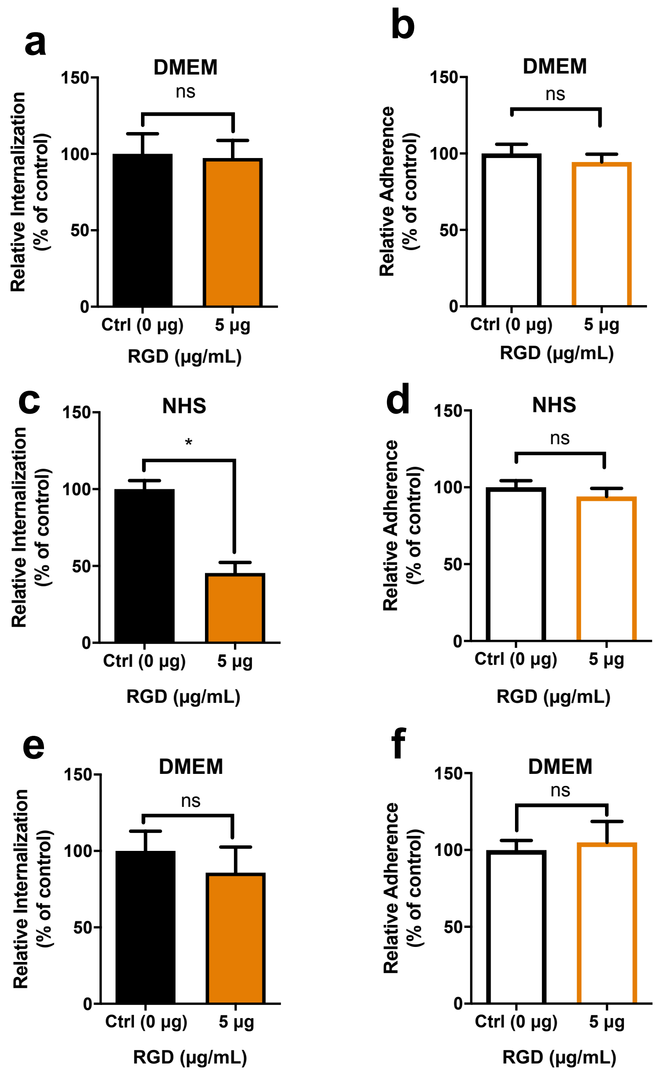

### Extended Data Fig. 9

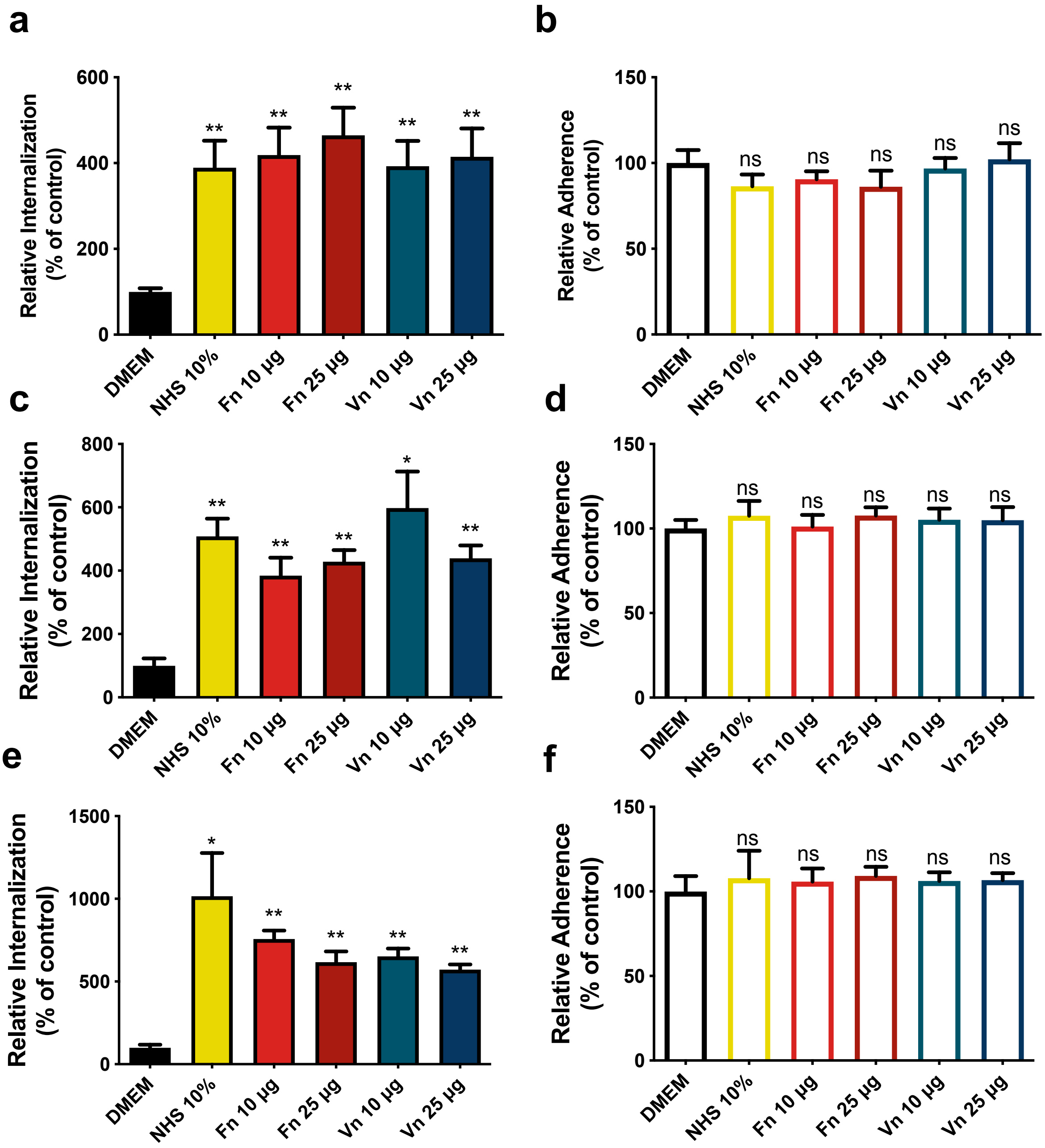

### Extended Data Fig. 10

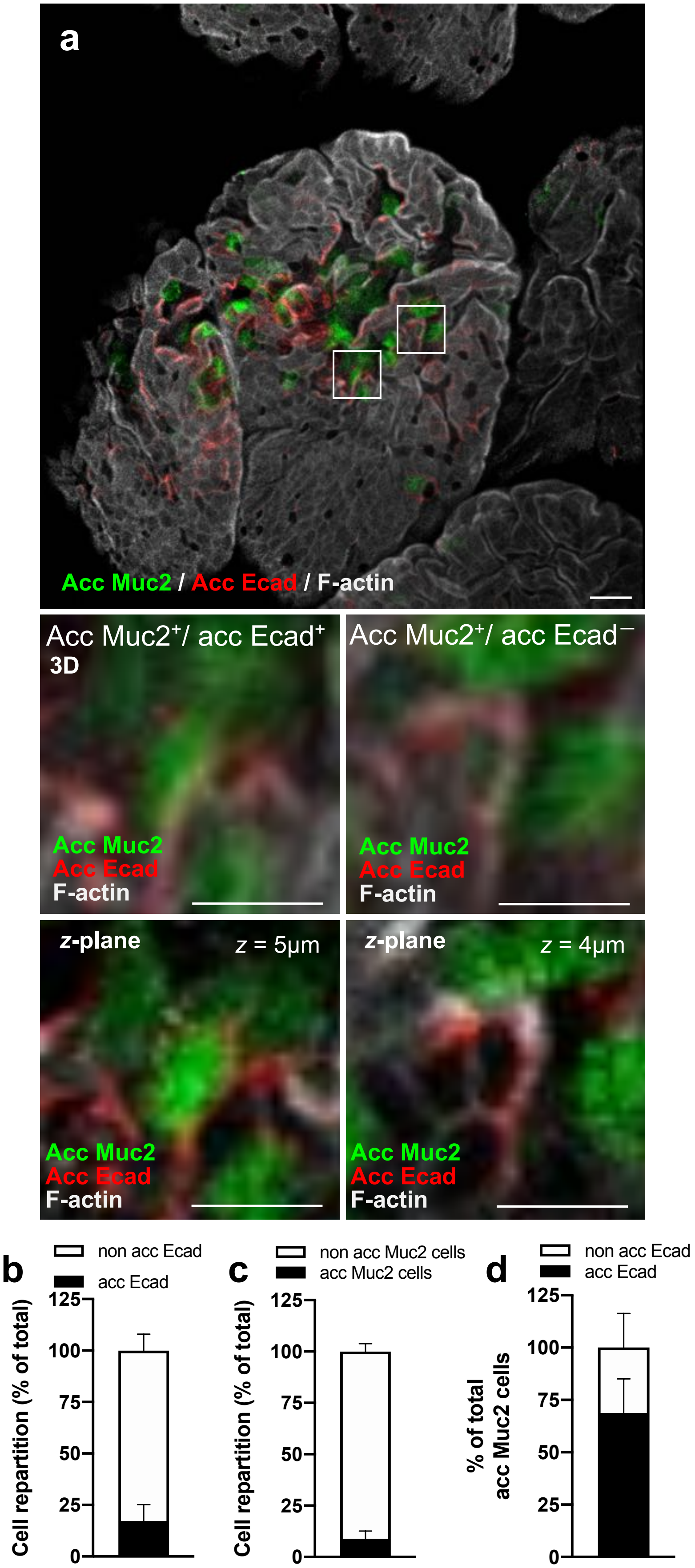

### Extended Data Fig. 11

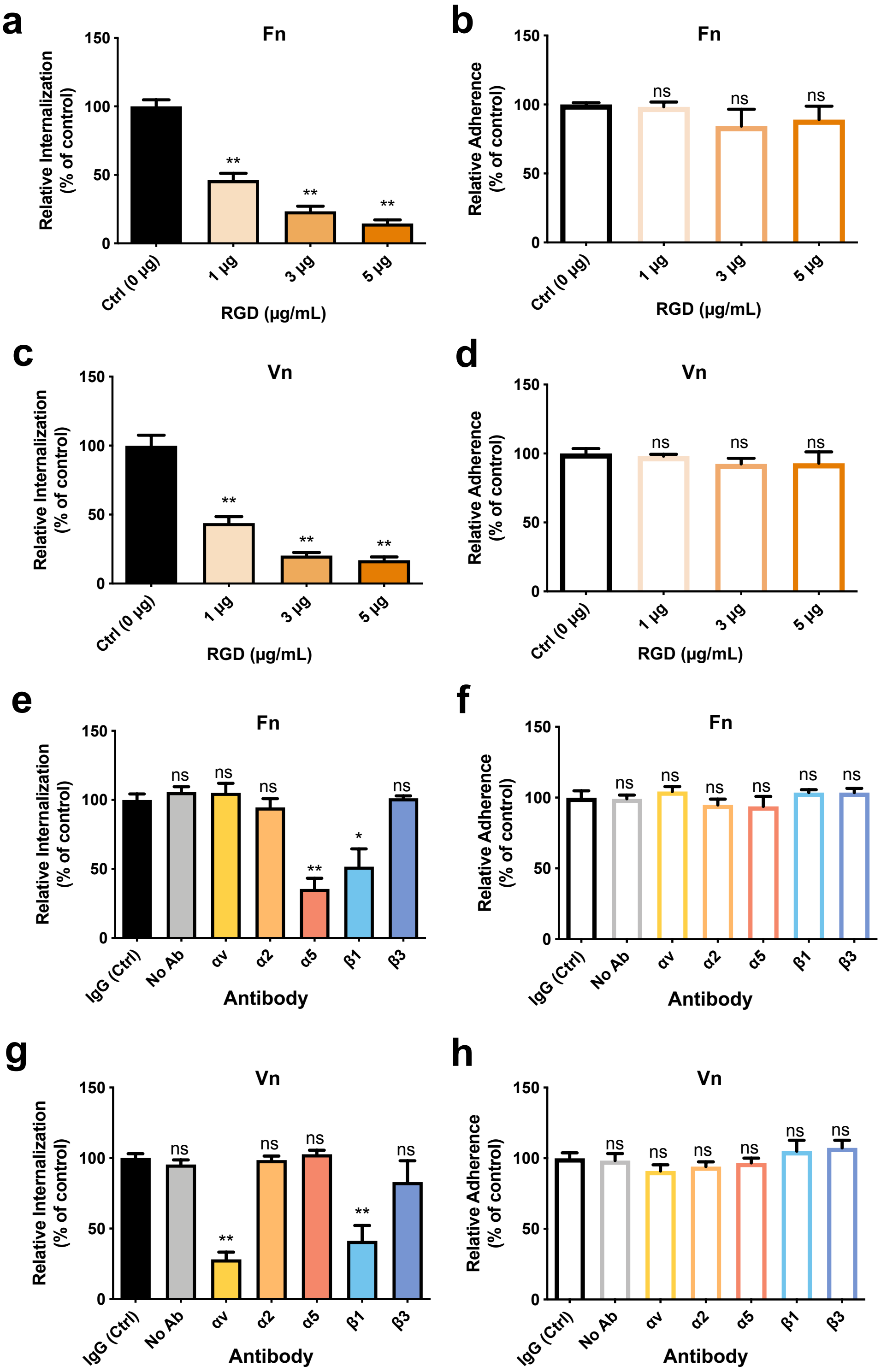

### Extended Data Fig. 12

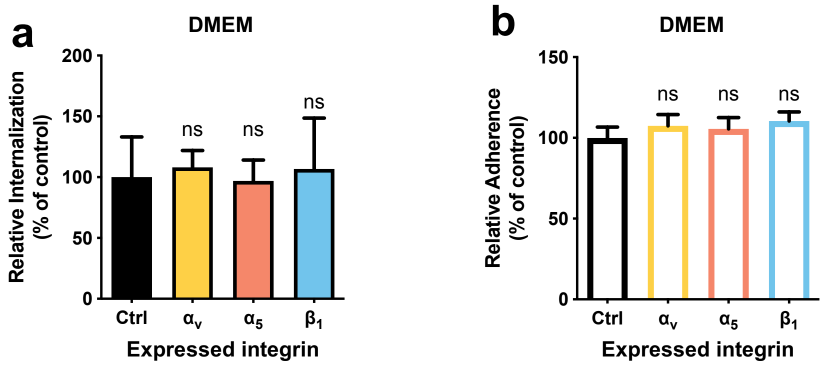

### Extended Data Fig. 13

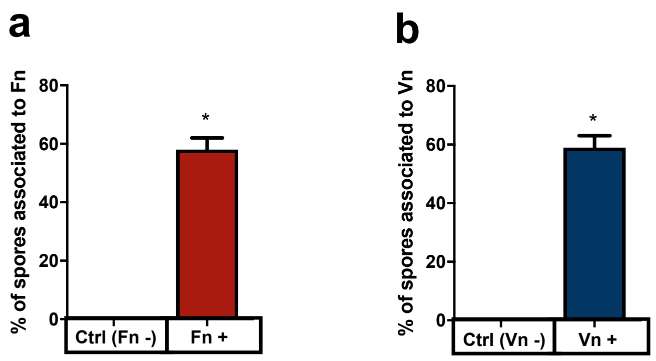

### Extended Data Fig. 14

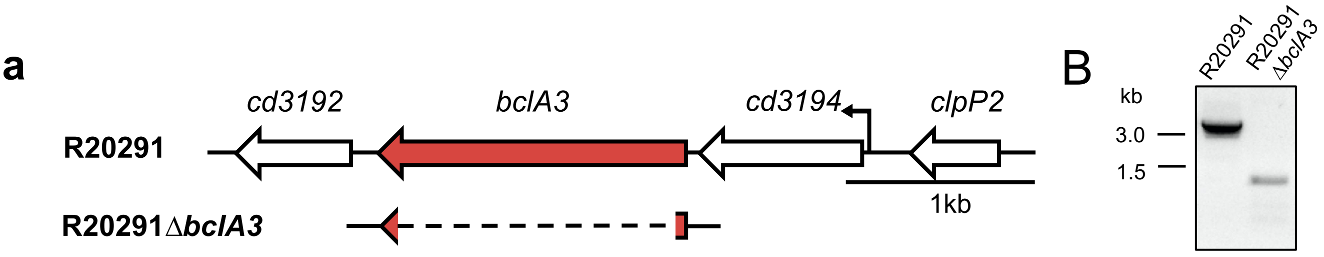

### Extended Data Fig. 15

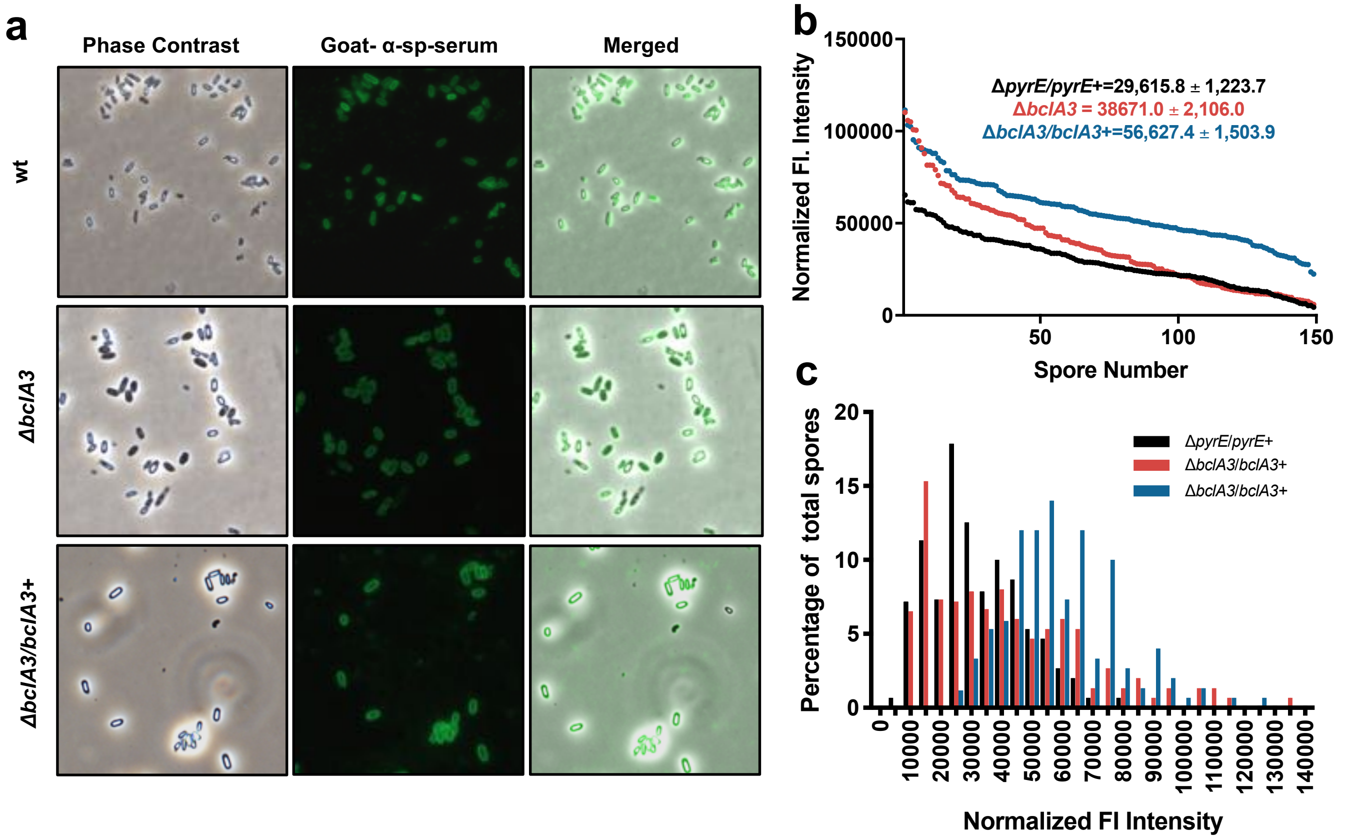

### Extended Data Fig. 16

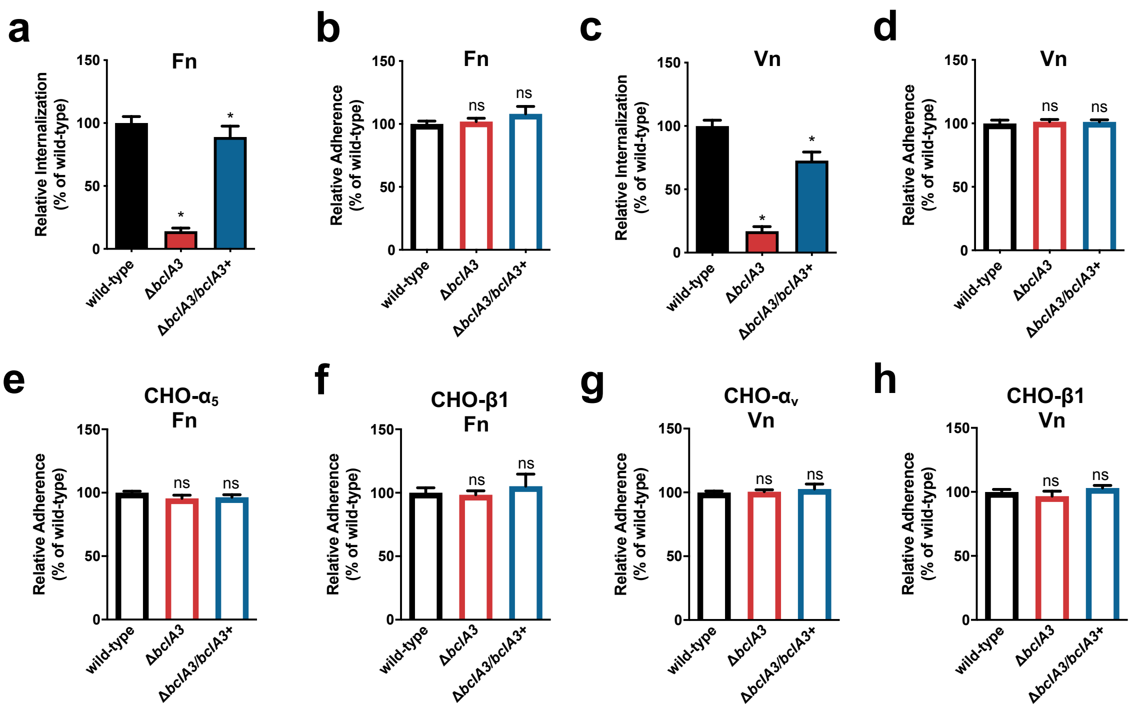

### Extended Data Fig. 17

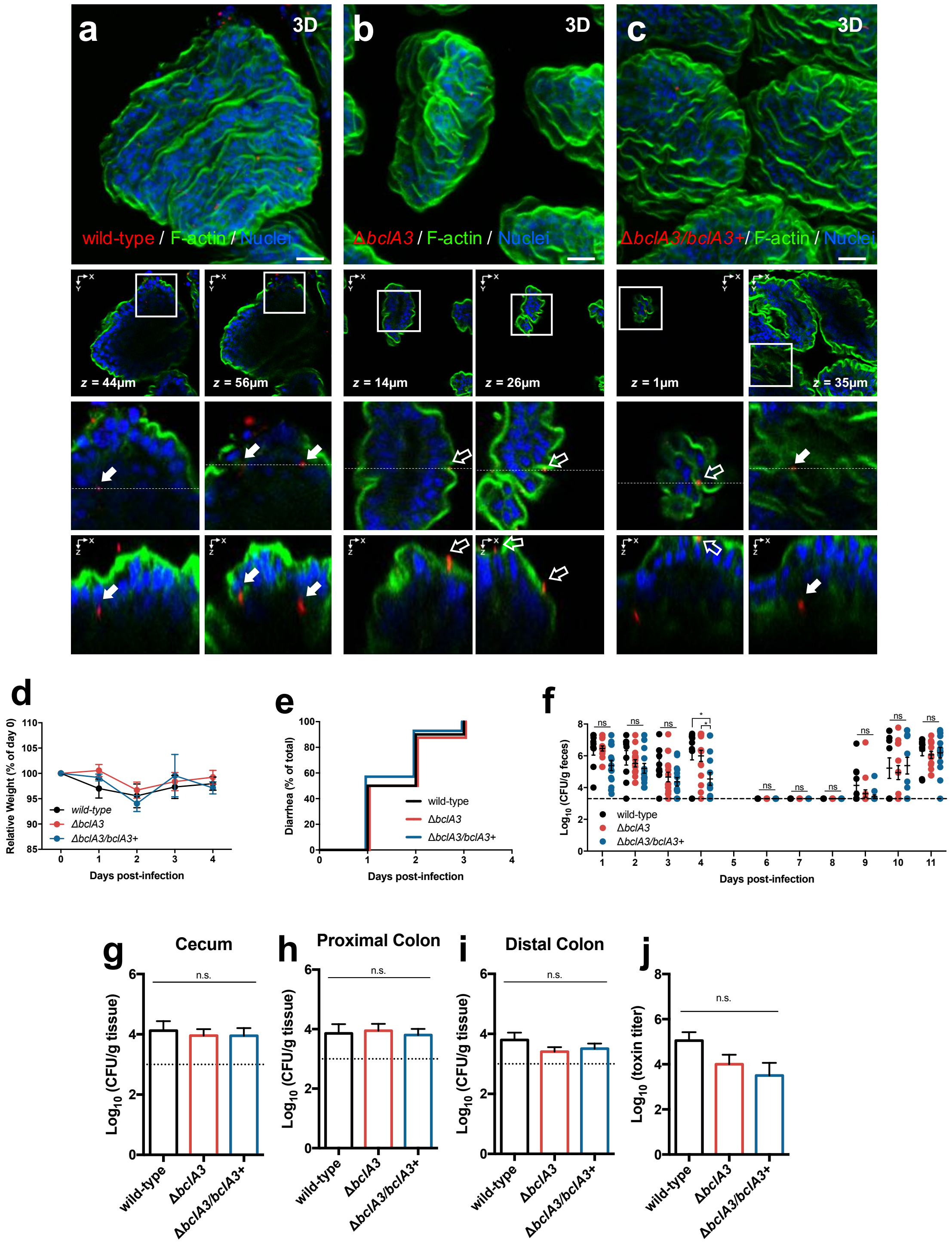

### Extended Data Fig. 18

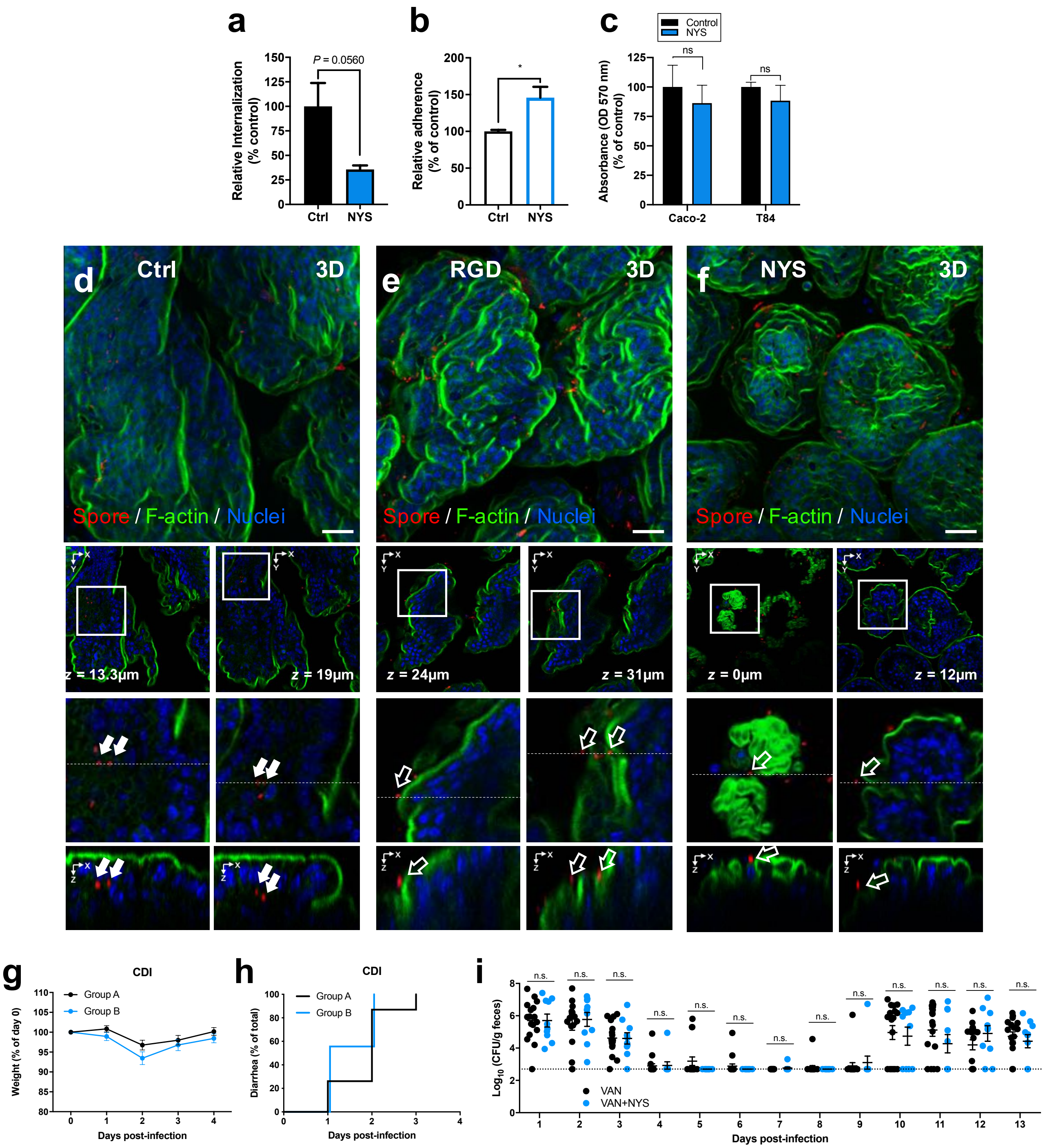
