## Supplementary Table S1 and S2 for "*Clostridioides difficile* spore-entry into intestinal epithelial cells contributes to recurrence of the disease"

**Supplemental Material**

**Supplementary Tables**

| TABLE S1. Bacterial strains and plasmids used | | |
| --- | --- | --- |
| Strain or Plasmid | Relevant characteristic | Source/Reference |
| *E. coli* Top10 | *mcrA* Δ(*mrr-hsdRMS-mrcBC*) *recA1* | Invitrogen |
| *E. coli* CA434 | *hsd20*(rB-, mB-, *recA13*, *rpsL20*, *leu*, *proA2*, with IncPb conjugative plasmid R702 | ^1^ |
| *C. difficile* 630Δ*erm* | A laboratory strain with erythromycin sensitive derivative of *C. difficile* strain 630 | ^2^ |
| *C. difficile* R20291 | Ribotype 027, epidemically relevant strain | ^3^ |
| *C. difficile* M120 | Ribotype 078 commonly isolated from farm animals | ^4^ |
| *C. difficile* PUC52 | Ribotype 027, isolated from a patient with CDI  Genome not available | ^5^^,^^6^ |
| *C. difficile* PUC30 | Ribotype 027, isolated from a patient with CDI  Genome not available | ^5^^,^^6^ |
| *C. difficile* PUC25 | Ribotype 014/020, isolated from a patient with CDI  Genome not available | ^5^^,^^6^ |
| *C. difficile* PUC31 | Ribotype 046, isolated from a patient with CDI  Genome not available | ^5^^,^^6^ |
| *C. difficile* PUC98 | Ribotype 082, isolated from a patient with CDI ^-^  Genome not available | ^5^^,^^6^ |
| *C. difficile* PUC131 | Ribotype 097, isolated from a patient with CDI ^-^  Genome not available | ^5^^,^^6^ |
| R20291 Δ*pyrE* | R20291 isogenic *pyrE* mutant | ^7^^,^^8^ |
| R20291 Δ*pyrE/pyrE+* | R20291 isogenic *pyrE* mutant complemented with wild-type *pyrE* into the *pyrE* loci | This work |
| R20291 Δ*bclA3* | R20291 isogenic *bclA3* mutant | This work |
| R20291 Δ*bclA3/bclA3+* | R20291 isogenic *bclA3* mutant complemented with wild-type *bclA3* in the *pyrE* loci | This work |
| ***Plasmids*** |  |  |
| pMTL-YN4 | Pseudo suicide vector containing: *Clostridium perfringens* *catP* resistant cassette, *Clostridium sporogenes* *pyrE;* Carries unaltered *colE1* *E. coli* replicon; *traJ* encoding transfer function of the RP4 *oriT* region; RepA and Orf2, the replication region of the *Clostridium botulinum* plasmid pBP1; and AscI/Sbf1 sites for the cloning of the right-hand homology arm/left-hand homology arm cassette. | ^7^^,^^8^ |
| pMTL-YN2C | Pseudo suicide vector containing: *Clostridium perfringens* *catP* cassette; *Clostridium sporogenes* *pyrE;* Carries unaltered *colE1* *E. coli* replicon; *traJ* encoding transfer function of the RP4 *oriT* region; RepA and Orf2, the replication region of the *Clostridium botulinum* plasmid pBP1; a left-hand homology arm encompassing a 300 bp internal fragment of the R20291 *pyrE* gene lacking 50 nucleotides from the 5´-end, and 235 bp from the 3´-end; a right-hand homology arm compromising the 1200 bp region of DNA immediately downstream of *pyrE*; this plasmid also has additional DNA segments inserted between the left-hand and right-hand homology arms which carries a copy of the *lacZ*´ containing a multiple cloning site region and a transcriptional terminator of the ferredoxin gene. | ^7^^,^^8^ |
| pDP376 | A 1086 bp PCR fragment was made by overlap extension PCR of a 544pb LHA upstream of the start codon of *bclA3* and a 542 bp RHA downstream of *bclA3* of strain R20291 The cassette was cloned by Gibson assembly into AscI/SbfI digested pMTL-YN4 vector. | This work |
| pMPG1 | A 3565-bp fragment containing the promoter region of the entire bicistronic operon formed by *sgtA* and *bclA3* of *C. difficile* strain R20291 was cloned into EcoRI and BamHI sites of pMTL-YN2C. Plasmid to complement the bclA3 mutant strain in the *pyrE* loci. | This work |

| Table S2. Primers used | | | | |
| --- | --- | --- | --- | --- |
| Primer code/Name | Primer sequence^a^ | Position^b^ | Gene | Use^c^ |
| P332  FP-LHA-bclA3- *pyrE* | *CCGATCGGGCCC*CCTGCAGGTGCTAGGTATAAAGAAGCAATAGAAGG | -544 to -517 | *bclA3* | MP |
| P334  RP-LHA-bclA3-pyrE | *CGATATTAGAAGCCTTTTCTTATCTAATCTA*TATTAAAAGCACCTCCTGATATATTTGAGC | -30 to -1 | *bclA3* | MP |
| P335  FP-RHA-bclA3-pyrE | *GCTCAAATATATCAGGAGGTGCTTTTAATA*TAGATTAGATAAGAAAAGGCTTCTAATATCG | +2034 to +2065 | *bclA3* | MP |
| P336  RP-RHA-b*clA3*-pyrE | *GCTAAGGATTCAGAAC*GGCGCGCCCCAGGAATAGCATATGAATTAGCCC | +2551 to +2576 | *bclA3* | MP |
| P664  FP-bclA3-detect | GAGAGGTATTTTATTCTGACATTGC | -699 to -673 | *bclA3* | MP |
| P665  RP-bclA3-detect | GCTTCCTATTCTTCCACAACACC | +2689 to +2712 | *bclA3* | MP |
| P476  NFP-bclA3c-promotor | *CCATGATTAC*GAATTCGAGGCTAAAGAGTACGGGCTGATTG | -1527 to -1502 | *bclA3* | CP |
| P477  NRP-bclA3c | *GACTCTAGA*GGATCCCTAATTTATTGCAATTCCTGCACTTGCATATCC | +2004 to +2037 | *bclA3* | CP |
| P530  FP-pyrE detect | AGAGAAGGAATAAAAAGTTTAGACGAAATAAGAGG |  |  | CP |
| P529  RP-pyrE detect | TTACATCCCTAATTCCTTGAACTCTC |  |  | CP |
| ^a^ Restriction sites are underlined; added sequence for proper restriction digestion are in italic; overlapping sequence for overlap extension PCR or Gibson cloning are in italic and grey.  ^b^ The nucleotide position numbering begins from the first codon and refers to the relevant position within the respective gene sequence. Restriction sites EcoRI (GAATTC), BamHI (GGATCC), Sbf1 (CCTGCAGG, AscI (GGCGCGCC) are underlined.  ^c^ MP, deletion of target gene; CP, complementation of target gene. | | | | |

**References**

1	Emerson, J. E. *et al.* A novel genetic switch controls phase variable expression of CwpV, a Clostridium difficile cell wall protein. *Mol Microbiol* **74**, 541-556.

2	Hussain, H. A., Roberts, A. P. & Mullany, P. Generation of an erythromycin-sensitive derivative of Clostridium difficile strain 630 (630Deltaerm) and demonstration that the conjugative transposon Tn916DeltaE enters the genome of this strain at multiple sites. *J Med Microbiol* **54**, 137-141.

3	McEllistrem, M. C., Carman, R. J., Gerding, D. N., Genheimer, C. W. & Zheng, L. A hospital outbreak of Clostridium difficile disease associated with isolates carrying binary toxin genes. *Clin Infect Dis* **40**, 265-272.

4	Goorhuis, A. *et al.* Emergence of Clostridium difficile infection due to a new hypervirulent strain, polymerase chain reaction ribotype 078. *Clin Infect Dis* **47**, 1162-1170.

5	Plaza-Garrido, A. *et al.* Predominance of *Clostridium difficile* ribotypes 012, 027 and 046 in a university hospital in Chile, 2012. *Epidemiol Infect* **144**, 976-979.

6	Plaza-Garrido, A. *et al.* Outcome of relapsing Clostridium difficile infections do not correlate with virulence-, spore- and vegetative cell-associated phenotypes. *Anaerobe* **36**, 30-38.

7	Ehsaan, M., Kuehne, S. A. & Minton, N. P. *Clostridium difficile* Genome Editing Using *pyrE* Alleles. *Methods Mol Biol* **1476**, 35-52.

8	Ng, Y. K. *et al.* Expanding the repertoire of gene tools for precise manipulation of the *Clostridium difficile* genome: allelic exchange using *pyrE* alleles. *PLoS One* **8**, e56051.
